# Cell-specific polymerization-driven biomolecular condensate formation fine-tunes root tissue morphogenesis

**DOI:** 10.1101/2024.04.02.587845

**Authors:** Jianbin Su, Xianjin Xu, Leland J. Cseke, Sean Whittier, Ruimei Zhou, Zhengzhi Zhang, Zackary Dietz, Kamal Singh, Bing Yang, Shi-You Chen, William Picking, Xiaoqin Zou, Walter Gassmann

**Affiliations:** Division of Plant Science and Technology; Interdisciplinary Plant Group, University of Missouri, USA; Dalton Cardiovascular Research Center; Department of Physics and Astronomy; Department of Biochemistry; Institute for Data Science and Informatics; Department of Veterinary Pathobiology; Department of Surgery; Christopher S. Bond Life Sciences Center, University of Missouri, USA; Donald Danforth Plant Science Center, St. Louis, Missouri, USA

**Keywords:** Biomolecular condensate, polymerization, lateral root cap (LRC), SRFR1, late embryogenesis abundant (LEA), dehydrin, DVL2

## Abstract

Formation of biomolecular condensates can be driven by weak multivalent interactions and emergent polymerization. However, the mechanism of polymerization-mediated condensate formation is less studied. We found lateral root cap cell (LRC)-specific SUPPRESSOR OF RPS4-RLD1 (SRFR1) condensates fine-tune primary root development. Polymerization of the SRFR1 N-terminal domain is required for both LRC condensate formation and optimal root growth. Surprisingly, the first intrinsically disordered region (IDR1) of SRFR1 can be functionally substituted by a specific group of intrinsically disordered proteins known as dehydrins. This finding facilitated the identification of functional segments in the IDR1 of SRFR1, a generalizable strategy to decode unknown IDRs. With this functional information we further improved root growth by modifying the SRFR1 condensation module, providing a strategy to improve plant growth and resilience.

## INTRODUCTION

Cells have evolved intricate signaling networks to respond to various developmental and environmental stimuli. Higher-order molecular assemblies play a key role in modulating such signaling characteristics such as specificity, magnitude, and duration (*1–3*). These higher-order assemblies can be classified as ordered supramolecular complexes with defined shapes formed by stable and comparatively rigid protein-protein interactions (signalosomes) or as labile oligomer assemblies with broad size distributions, ranging from small clusters composed of several molecules to large biomolecular condensates that provide an alternative strategy for temporal and spatial organization of cellular constituents (*4–10*). Moreover, some biomolecular condensates are dynamic in size and composition, making biomolecular condensates ideally suited for fine-tuning cellular signaling dynamics (*11–14*).

Multivalency, the driving force for biomolecular condensate formation, can be provided by either intrinsic sticker-sticker interactions encoded in intrinsically disordered regions (IDRs) or by emergent polymerization of folded domains (*9, 10*). IDRs and intrinsically disordered proteins (IDPs) are defined as polypeptide chains that lack stable tertiary structures (*15, 16*). The specific combinations of sticker binding properties, such as hydrogen bonding, ionic interactions, cation-p, p-p, and hydrophobic interactions, give rise to different thermodynamic and kinetic properties of biomolecular condensates (*9*). Less studied are the principles of polymerization-mediated biomolecular condensate formation (*9, 17–20*). Considerable studies have focused on IDRs as the drivers for biomolecular condensate formation. A tuner function of IDR has also been found for condensates that are mediated by oligomerization and aggregation-prone domains (*21–24*). It is likely that a generalized IDR mechanism of fine-tuning spontaneous aggregation may account for a considerable portion of biomolecular condensate formation. Yet, the mechanisms of how IDRs tune the polymerization or aggregation properties of folded domains are largely unknown.

Protection of enzymes from irreversible aggregation has been extensively studied for plant late embryogenesis abundant (LEA) proteins, a large family of IDPs (*25*). In addition to stochastic environmental drought stress, plant seeds undergo developmentally programmed desiccation during maturation. Accumulation of LEA proteins has been found to play a key role in protecting enzymes from aberrant aggregation during environmental dehydration and developmental desiccation (*25–30*). At the molecular level, the protective function of LEA proteins is mainly mediated by molecular shielding effected by non-specific transient interactions with their target proteins, through which aggregation can be avoided (*25, 30*). Due to the functional similarity between LEA proteins and IDRs that function as tuners for aggregation-prone domains, we explored whether information on IPDs can serve as a guide to decode the molecular mechanisms of IDRs in biomolecular condensate formation.

In this study, we found that the Arabidopsis SRFR1 protein specifically forms biomolecular condensates in lateral root cap cells (LRCs) and that these condensates are required for optimal root growth under varying growth conditions. Interestingly, three plant dehydrins, a class of IDPs that protect cellular enzymes from aggregation during dehydration and desiccation (*25, 26, 30–32*), can functionally substitute for IDR1^SRFR1^. Informed by the dehydrin mode of action, we further identified several segments in IDR1^SRFR1^ that are essential for tuning its adjacent polymerizing domain during biomolecular condensate formation. Manipulation of these segments resulted in engineered plants with reduced fluidity of SRFR1 condensates and longer primary roots. Our study demonstrated a quantitative relationship between parameters of biomolecular condensate formation and plant growth, offering a strategy to improve resilience in fluctuating environments.

## RESULTS

### SRFR1 regulates primary root growth

In our previous work, we demonstrated that SUPPRESSOR of rps4-RLD 1 (SRFR1) is a negative regulator of effector-triggered immunity (ETI) mediated by several nucleotide-binding domain leucine-rich repeat containing receptors (NLRs), including SNC1, RPS6, RPS4-RRS1 and RPS4B-RRS1B (*33–35*). As a result of the variation in NLR family members between accessions, *srfr1* mutants exhibited different phenotypes depending on the genetic background (fig. S1, A-D). For example, *srfr1-4* exhibits severe stunting which is mediated by the Col-0 specific NLR protein SNC1 (fig. S1, B and E). The Wassilewskija-0 (Ws-0) background CRISPR mutant *srfr1^WS^* grew normally during the first three weeks, then cell death began to expand from petiole to leaf tip (fig. S1D). Apart from *SNC1*-mediated shoot stunting in Col-0 we observed a modest reduction of shoot growth in RLD and Ws-0 *srfr1* mutants (fig. S1, C and D) (*34*).

Interestingly, all *srfr1* mutants, including newly CRISPR-generated alleles (fig. S2A), exhibited a shortened primary root (fig. S1, F-H, and S2B). The accession-independent short primary root phenotype of *srfr1* mutants suggested that regulating primary root growth might be a general function of SRFR1 and might be unrelated to ETI. Indeed, *srfr1-4 snc1-11* and *srfr1-4 eds1-2* still exhibited short primary roots, in contrast to the observed reversion of shoot stunting by mutations in *EDS1* or *SNC1* (fig. S1, E and F) (*35, 36*). Next, we asked whether the short primary root phenotype of *srfr1* mutants is caused by elevated stress responses. Because ethylene acts as a key regulator of stress-induced morphological responses in roots (*37*), we first treated *srfr1-1* and *srfr1-2* with AgNO3 and aminoethoxyvinylglycine (AVG), two ethylene biosynthesis inhibitors. As expected, the short root phenotype of *srfr1-1* and *srfr1-2* was rescued by AgNO3 and AVG (fig. S2C). Consistent with this observation, several *ACS* genes encoding enzymes that catalyze the rate limiting step of ethylene biosynthesis, mainly *ACS7* and *ACS9*, were expressed more highly in root tissue of *srfr1* mutants (fig. S2D). To provide genetic evidence that the short primary root phenotype results from elevated ethylene responses, we crossed *srfr1-4* with *etr1-3* and *ein2*, two key ethylene signaling mutants. As shown in fig. S1G, the short root phenotype of *srfr1-4* was rescued completely by mutations in ethylene signaling. Together, these results suggest that regulation of primary root growth by SRFR1 is dependent on ethylene signaling and that SRFR1 is required to avoid overactivating ethylene responses.

### Lateral root cap SRFR1 condensate formation correlates with optimal root growth

To explore the molecular mechanism of primary root growth regulation by SRFR1, we first investigated the developmental basis for this phenotype. Most prominently, the shorter root of *srfr1* mutants correlated with a reduced root meristem size (fig. S1, I and J). We next examined the tissue and subcellular localization of SRFR1 in roots. We inserted a YFP and an HA tag just before the start codon in a 9 kb *SRFR1* genomic clone, designated *YFP-HA-gSRFR1^WT^*. This construct complemented all mutant phenotypes of *srfr1-1* and *srfr1-4* (Fig. 2D-F and fig. S6), suggesting that the function of SRFR1 is not affected by the N-terminal YFP-HA tag. SRFR1 mainly displayed a diffuse cytoplasmic distribution in most root tissues but exhibited punctate localizations in upper lateral root cap cells (LRCs) (Fig. 1, A and B, and fig. S3A). More precisely, SRFR1 puncta were found only in upper LCRs (Fig. 1A). In shoot tissues, SRFR1 puncta were frequently found in developing trichomes and young stomatal cells (fig. S3B). In contrast, SRFR1 puncta were rarely observed in mesophyll cells (fig. S3B). Collectively, these results suggest that SRFR1 puncta represent a spatially and temporally distinct tissue-specific biomolecular condensate.

**Fig. 1.**
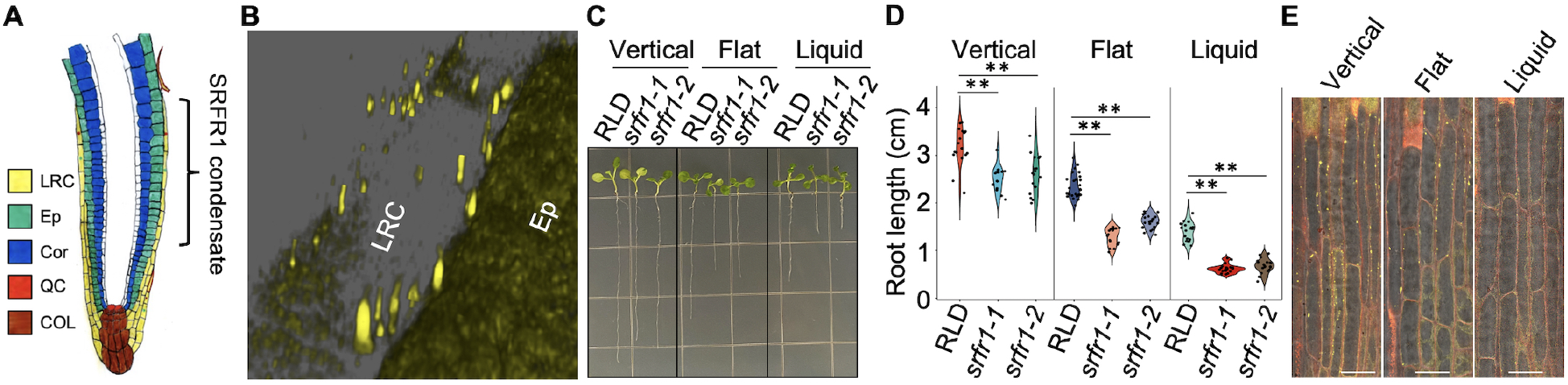
Lateral root cap SRFR1 condensate is responsive to growth conditions. (**A**) Schematic diagram of Arabidopsis root apical meristem localization of SRFR1 condensate. LRC: lateral root cap; Ep: epidermis; Cor: cortex; QC: quiescent center; COL: columella. (**B**) Subcellular localization of SRFR1 in the lateral root cap and epidermis. Z-stack of 6-day-old seedling roots were taken with confocal microscopy. (**C** and **D**) Primary root length of *srfr1-1* and *srfr1-2* are sensitive to growth conditions. Primary root length is measured with 8-day-old seedlings, n = 14-25. ** indicates p < 0.001. (**E**) Accumulation of LRC SRFR1 condensates are responsive to growth conditions. Single median section images were taken with roots of 6-day-old *srfr1-1 YFP-HA-gSRFR1^WT^* #7 seedlings grown under different conditions. Bar = 25 μm.

**Fig. 2.**
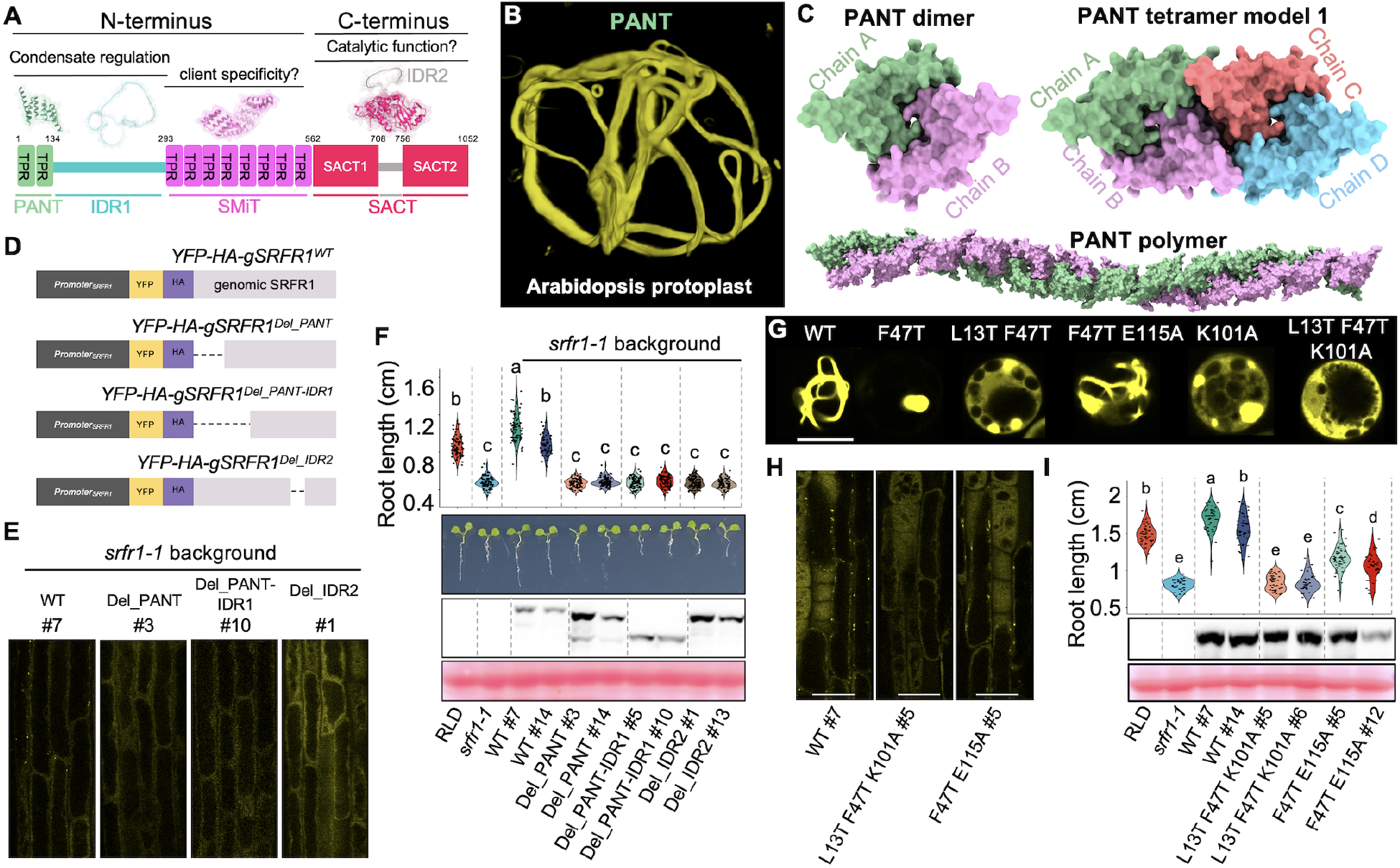
PANT domain polymerization is required for lateral root cap SRFR1 biomolecular condensate formation and primary root growth. (**A**) Schematic diagram of SRFR1 protein architecture. (**B**) 3-D localization of PANT domain. (**C**) PANT dimer and tetramer predicted are by AlphaFold2. The PANT 24-mer is generated using MatchMaker. (**D**) Schematic diagram of full-length and truncated SRFR1 constructs used for genetic complementation assays. (**E**) Subcellular localization of full-length and truncated SRFR1 in lateral root cap. (**F**) Deletion of PANT, PANT-IDR, or IDR2 failed to rescue the *srfr1-1* short primary root phenotype. Primary root length is measured with 6-day-old seedlings, n = 63-88. Ponceau S staining of the Rubisco large subunit shows equal protein loading. (**G**) Subcellular localization of PANT dimer and tetramer interface variants in rice protoplasts. Bar = 10 μm. (**H**) Subcellular localization of YFP-HA-tagged SRFR1 dimer- and tetramer-interface variants in the lateral root cap. Bar = 10 μm. (**I**) Primary root length and protein expression of YFP-HA-tagged SRFR1 dimer- and tetramer-interface variants. Primary root length is measured with 6-day-old seedlings, n = 32-51. Ponceau S staining of the Rubisco large subunit shows equal protein loading. Violin-plots show the data distribution. Letters denote different statistical groups (ANOVA, Tukey-Kramer grouping).

Plant root growth is highly plastic and is determined by internal developmental cues as well as by integration of environmental signals. In a first limited test of growth conditions, we found that root growth of *srfr1* mutants was less affected (∼ 19.5% reduction) when seedlings were grown vertically on ½ MS agar plates with whole roots exposed to air, compared to when they were grown horizontally (∼ 39% reduction) with roots penetrating the growth media. Root length reduction was even stronger when seedlings were grown in liquid ½ MS medium (54.6% reduction) (Fig. 1, C and D). Thus, we asked whether SRFR1 condensates were differentially regulated under these conditions. We found a clear correlation between SRFR1 condensate formation and primary root length (Fig. 1E). Further, SRFR1 condensates were completely eliminated when roots were treated with ACC, BAP, and IAA, known inhibitors of root growth (fig. S4). AVG and IBA application had weaker effects (fig. S4). Because IBA needs to be converted to IAA in LRCs for its function (*38*), it is likely that its conversion is less efficient under our experimental conditions. The dynamic accumulation of SRFR1 condensates suggests a role for SRFR1 in facilitating optimal root growth in response to environmental and developmental stimuli.

### The PANT-IDR1 module is required for LRC SRFR1 biomolecular condensate formation and optimal primary root growth

Structural prediction by Alphafold2 suggests that SRFR1 is a protein with 3 distinct domains and 2 intrinsically disordered regions (IDRs) (Fig. 2A). We named the domains plant-associated N-terminal TPR (PANT) domain, SRFR1 middle TPR (SMiT) domain, and SRFR1-associated C-terminal domain (SACT). IDR1 separates the PANT and SMiT domains, and the shorter IDR2 separates the SACT domain into two subdomains (Fig. 2A). The PANT domain is conserved in plants but absent in animals. The SACT domain is conserved in eukaryotes and always associated with at least a SMiT domain, but its function is unknown (*33*). To ask which domain is required for SRFR1 biomolecular condensate formation, we expressed each region and their combinations in both rice and Arabidopsis protoplasts, a heterologous and homologous system, respectively. Some combinations were also tested in human HEK293T cells. Full-length SRFR1 formed small puncta in both rice and Arabidopsis protoplasts (fig. S5, A and C). Surprisingly, the PANT domain alone formed cytoskeleton-like fibrils in Arabidopsis, rice, and human cells (Fig. 2B, fig. S5 A-E). 3-D reconstruction showed that PANT fibrils extended throughout cells (Fig. 2B, fig. S5B, and movie. S1).

Interestingly, PANT fibril formation was suppressed in the presence of IDR1, resulting in only one or two puncta in rice protoplasts but diffuse localization in Arabidopsis protoplasts and human HEK293T cells (fig. S5A-D). Unlike prion-like IDRs that promote multivalent weak interactions and drive biomolecular condensate formation (*9*), our results suggest that IDR1 of SRFR1 is a polymerization-suppressing IDR. Combined, we hypothesize that SRFR1 condensate formation is driven by PANT polymerization and fine-tuned by IDR1. This hypothesis is supported by the result that SRFR1 condensates were completely abolished when the PANT domain was removed from otherwise full-length SRFR1, while we still observed puncta when IDR1 or IDR2 were deleted (fig. S5, B and C).

We next verified these results from transient overexpression using stable transgenic plants. PANT, PANT-IDR1, or IDR2 were deleted in the native promoter-driven genomic clone *YFP-HA-gSRFR1^WT^* (Fig. 2D). These constructs were used to transform both *srfr1-1* and *srfr1-4*.

Deletion of either PANT, PANT-IDR1 or IDR2 resulted in high protein accumulation in both *srfr1-1* and *srfr1-4* (Fig. 2F and fig. S6B). However, SRFR1 with deleted PANT, PANT-IDR1 or IDR2 failed to complement the short primary root phenotype and abolished LRC SRFR1 biomolecular condensate formation despite high protein levels (Fig. 2, E and F). To our surprise, the PANT and IDR1 domains were dispensable for complementation of ETI-related phenotypes, such as severely stunted shoots of *srfr1-4* (fig. S6), suggesting that PANT-IDR1-mediated SRFR1 biomolecular condensate formation is not required for suppressing ETI. However, SRFR1 without PANT or PANT-IDR1 was not as efficient in promoting wild type-level shoot growth (fig. S6), suggesting that dynamic condensate regulation mediated by PANT-IDR1 also plays a role in buffering daily environmental fluctuations for optimal shoot growth. Finally, the short 51 aa IDR2 (708–758) is essential, since SRFR1 with IDR2 deleted failed to complement all *srfr1* mutant phenotypes (Fig. 2, E and F). In summary, transient expression in protoplasts and stable expression in the whole organism support the PANT-IDR1 module is required for LRC SRFR1 biomolecular condensate formation and optimal growth.

### Function of LRC SRFR1 biomolecular condensates requires polymerization of the PANT domain

The fibril-like localization pattern suggests PANT is capable of spontaneous polymerization. To gain insights into PANT polymerization, we modeled the structures of PANT dimers and tetramers using AlphaFold2 (Fig. 2C) (*39*). The top-ranked structure by AlphaFold2 suggests that PANT dimers are stabilized by multiple interactions, including the salt bridge between K111 and E115, hydrogen bonds between S18 and E117 and between Q74 and D109, as well as hydrophobic interactions contributed by F11, C17, W22, F47, L104, V110, and L114 (movie. S2). This model was then validated by mutational analysis (Fig. 2G, fig. S7, A and B). In rice protoplasts, the localization pattern of PANT variants could generally be classified into four categories: 1) WT-like fibrils (e.g., L13T); 2) short or thin fibrils (e.g., E115A); 3) puncta and puncta with emerging thin fibrils (e.g., F47T); and 4) completely diffuse (e.g. F47T/V110T). Collectively, these results suggest that hydrophobic interactions play a significant role in PANT dimer formation, since F47T/V110T or V110T/L114T double mutations were sufficient to cause entirely diffuse localization (fig. S7A). It is worth noting that the localization of some PANT variants differed significantly in rice and Arabidopsis protoplasts. The PANT variants F47T, V110T, K111A, and L114T formed ring-like structures around chloroplasts in Arabidopsis protoplasts (fig. S7B), suggesting that the PANT domain is regulated by additional factors in its native system, such as a co-evolved chaperone network.

According to AlphaFold2 predictions, PANT tetramers may exist as ensembles of several structural conformations (Fig. 2C, fig. S7D, and movie S2). Among these, model 1 is highly represented (11 out of 20 predicted models) with both electrostatic interactions mediated by E12- K15-Q90 and E97-R124-Q98-K101, and hydrophobic interactions mediated by F11-L94-L104. This model was supported by the punctate and diffuse localization of mutations K15A, K101A, and R124A (fig. S7C). Based on model 1, a 24-mer PANT linear polymer was built (Fig. 2C and movie. S2). Notably, mutations K15A, K101A, and R124A did not result in complete diffused localization (fig. S7C), suggesting that model 2 and model 3 may also represent PANT polymerization under certain physiological conditions (fig. S7D).

Next, we tested whether PANT polymerization is required for LRC condensate formation and primary root growth regulation. To ensure a complete abolishment of SRFR1 condensate formation, we generated a SRFR1^L13T^ ^F47T^ ^K101A^ variant with mutations in both PANT dimer and tetramer interfaces. Diffuse localization of PANT^L13T^ ^F47T^ ^K101A^ was first confirmed in rice protoplasts (Fig. 2G and fig. S7C). We also generated a SRFR1^F47T^ ^E115A^ variant for genetic complementation assays, since PANT^F47T^ ^E115A^ produced short and fragmented fibrils (Fig. 2G and fig. S7A). As expected, SRFR1^L13T^ ^F47T^ ^K101A^ exhibited diffuse localizations in LRCs and failed to complement the short root phenotype of *srfr1-1* (Fig. 2H and I). In addition, SRFR1^F47T^ ^E115A^ produced fewer condensates in LRCs and resulted in intermediate primary root growth (Fig. 2H and I). Our results support a quantitative positive relationship between PANT polymerization mediated SRFR1 biomolecular condensate formation in LRCs and primary root growth.

### IDR1 of SRFR1 can be functionally substituted by plant dehydrins

Our subcellular localization assays suggested that PANT polymerization drives, while IDR1 fine-tunes, SRFR1 condensate formation. Next, we asked which features of IDR1^SRFR1^ determine its functionality. Interestingly, we found PANT-IDR1^SRFR1^ exhibited completely diffuse localization in rice protoplasts when a 6 aa tail (PGIHLI) was added by the multiple cloning site of *pSAT6-eYFP* construct. We then took advantage of this localization pattern to explore substitutions that mimic the function of native IDR1^SRFR1^ (fig. S8A). First, we substituted Arabidopsis IDR1^SRFR1^ with the IDR1 from its orthologs. PANT-IDR1^GmSRFR1^ (soybean) and PANT-IDR1^OsIDR1^ (rice) exhibited diffuse localization, whereas formation of puncta was observed for PANT-IDR1^NtSRFR1^ (tobacco) and PANT-IDR1^ZmSRFR1^ (maize) (fig. S8, A and B). We next substituted IDR1^SRFR1^ with intrinsically disordered proteins (IDPs) or IDRs with well-studied features, including 7 plant late embryogenesis abundant (LEA) proteins and 4 prion-like IDRs from plant and human proteins. As shown in fig. S8A, PANT fusion with 4 of the 5 group II LEA proteins (also known as dehydrins) resulted in diffuse protein localization, including HIRD11, XERO1, XERO2, and COR47, mimicking IDR1^SRFR1^ function. Interestingly, PANT fusion with ERD14 resulted in large vesicle-like structures, some of which were as large as chloroplasts. Fusion of PANT with COR15A or LEA4-1, belonging to type III and type IV LEA proteins, respectively, caused aggregation-like punctate localization. Plant dehydrins, which are composed of conserved segments such as K-, Y-, S-, and F-segments separated by flexible linkers(*25, 26, 30–32*), prevent protein aggregation during developmental and abiotic dehydration processes (*25, 27–29, 32, 40*). Conceptually, it is therefore likely that IDR^SRFR1^ functionally mimics the mode of action of dehydrins by inhibiting the polymerization of the PANT domain.

In contrast to the function of dehydrins, multivalent weak interactions contributed by prion-like IDRs, also known as prion-like domains (PrLD), drive biomolecular condensate formation (*9, 10*). Fusion of PANT with 3 of the 4 prion-like IDRs exhibited punctate or aggregation-like localizations (fig. S8, A and B) (*41–44*), confirming the distinct functionality of dehydrins and PrLDs and that IDR1^SRFR1^ mimics the mode-of-action of dehydrins to fine-tune PANT condensation.

Next, we substituted HIRD11, ERD14 and COR47 for IDR1^SRFR1^ in the genomic clone *YFP-HA-gSRFR1^WT^*, designated as *YFP-HA-gSRFR1^IDR1-HIRD11^*, *YFP-HA-gSRFR1^IDR1-ERD14^*, and *YFP-HA-gSRFR1^IDR1-COR47^*, respectively (Fig. 3A). To our surprise, all these dehydrin substitutions rescued the short root phenotype of *srfr1-1* (Fig. 3B and fig. S8C). Strikingly, SRFR1^IDR1-ERD14^ even produced significantly longer roots (Fig. 3B and fig. S8C), possibly due to higher protein accumulation.

**Fig. 3.**
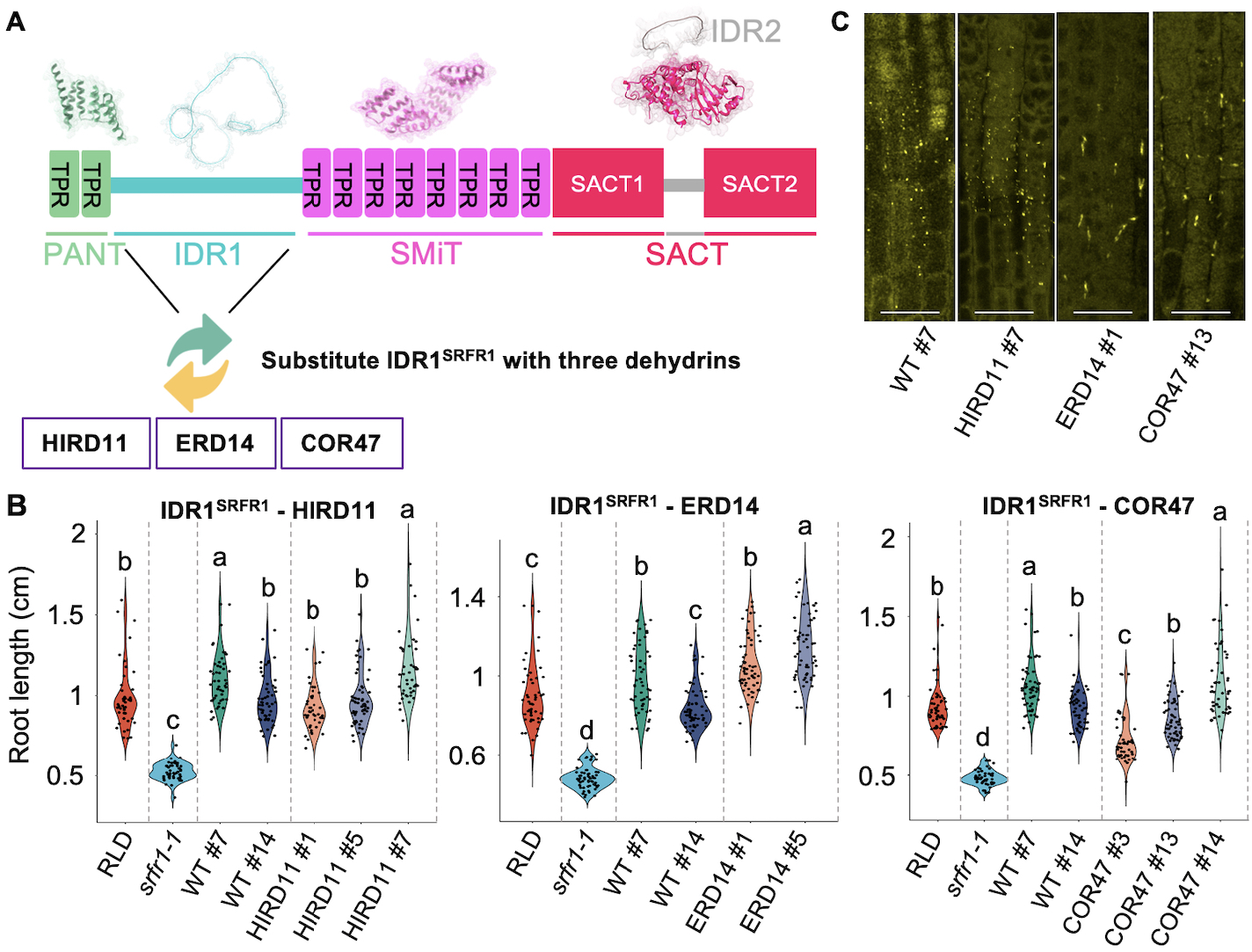
Group II LEA functionally substitutes for IDR1^SRFR1^. (**A**) Schematic diagram of functional substitution strategy. (**B**) Primary root length of YFP-HA-tagged SRFR1 with HIRD11, ERD14 or COR47 IRD1 substitutions. Primary root length is measured with 6-day-old seedlings, n = 42-65. Violin-plots show the data distribution. Letters denote different statistical groups (ANOVA, Tukey-Kramer grouping). (**C**) Lateral root cap biomolecular condensate of YFP-HA-tagged SRFR1 with HIRD11, ERD14 or COR47 IDR1 substitutions. Maximal projection images are shown, bar = 25 μm.

We also examined the formation of biomolecular condensates by dehydrin substitution constructs in LRCs (Fig. 3C). SRFR1^IDR1-HIRD11^ formed condensates similar in morphology to wild-type (WT) SRFR1, while SRFR1^IDR1-ERD14^ and SRFR1^IDR1-COR47^ substitutions formed significantly larger condensates. The protein abundance of some dehydrin substitution lines was significantly higher than WT, yet no alteration in growth was observed for these plants. Taken together, these results suggest that SRFR1 condensate formation in LRCs promotes root growth with a low minimum required amount and a buffered maximum. These observations prompted us to explore functional segments within IDR1^SRFR1^ based on the known dehydrin modes of action.

### IDR1^SRFR1^ fine-tunes SRFR1 condensate formation and primary root growth

Currently, predicting binding of disordered regions to folded domains remains challenging (*45*). To examine potentially labile secondary structures in IDR1^SRFR1^, we first divided it into short segments when encountering proline, a helix disruptor, as well as serine and glycine, which have moderate-to-high rotational freedom and are frequently found in random coils. Secondary structures of these segments were then predicted with PEP-FOLD3.5 (*46*), an online tool for *de novo* peptide structure prediction. Based on these principles, we identified 5 segments that might form labile secondary structures, namely P1-P5 (Fig. 4A and fig. S9A). To gain insights into the regulatory mechanism of IDR1^SRFR1^, all 5 potential segments were included in our mutational analysis.

**Fig. 4.**
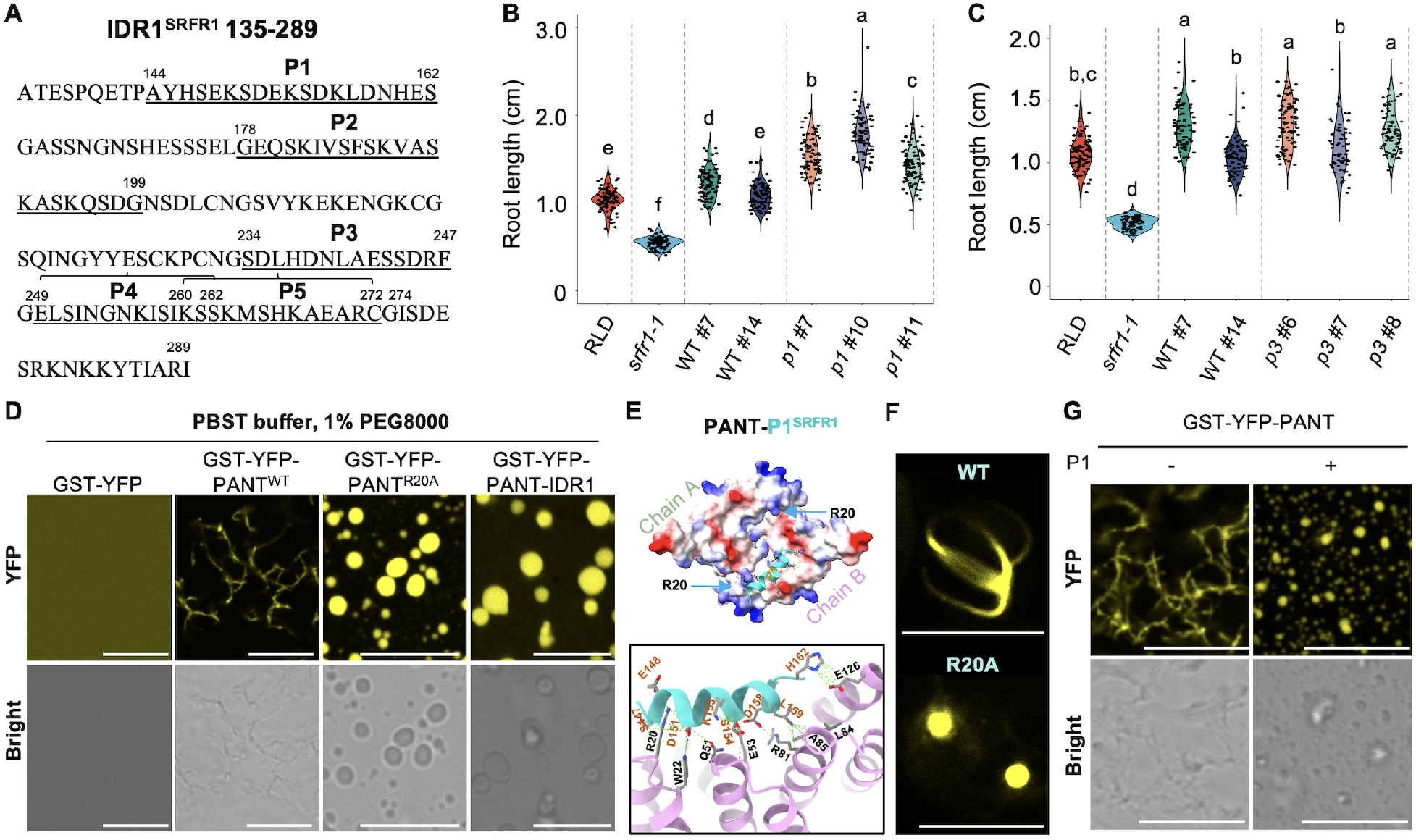
IDR1^SRFR1^ tunes SRFR1 biomolecular condensate formation and primary root growth. (**A**) Potential functional segments in IDR1^SRFR1^. (**B** and **C**) Primary root length of YFP-HA-tagged SRFR1 *p1* (E148K E152K D155N D158N) and *p3* (D235H D238H E242H D245K) variants. Primary root length is measured with 6-day-old seedlings, n = 58-93. Violin-plots show the data distribution. Letters denote different statistical groups (ANOVA, Tukey-Kramer grouping). (**D**) Fibril and condensate formation by PANT, PANT^R20A^, and PANT-IDR1^SRFR1^. Images were taken with the same confocal microscopy settings except for panels without fibril or condensate formation which were imaged with higher laser power and gain. Bar = 10 μm. (**E**) Interaction between PANT dimer and P1 peptide predicted by MDockPeP2. (**F**) Subcellular localization of PANT variants in rice protoplasts. Bar = 10 μm. (**G**) The P1 peptide inhibits PANT fibril formation and promotes spherical droplet formation. Bar = 10 μm.

P1 was predicted to form a helix with its positively and negatively charged surfaces facing opposite sides. As shown in fig. S9A, double mutations in P1 (E148K/E152K) resulted in both dispersed and punctate localizations, while quadruple mutations of P1 (E148K/E152K/D155N/D158N) shifted protein localization to more irregularly-shaped aggregation-like puncta. P3 is predicted to be an amphipathic helix with a hydrophobic and a negatively charged surface. Similar to P1, the double-mutation D235H/D238H in the negatively charged surface resulted in both dispersed and punctate localization, while the quadruple mutation D235H/D238H/E242H/D245K shifted the localization to nearly complete aggregate-like structures, indicating less fluidity. Mutations L236S/L240S in the hydrophobic surface showed no effect.

To probe inter-segment interactions, we performed N-terminal and C-terminal truncations in IDR1^SRFR1^. As shown in fig. S9A, the N-terminal part of IDR1^SRFR1^ (135–213) covering P1 and P2 is sufficient to suppress PANT condensation. Because the acidic amino acids in P1 and P3 are important for PANT-IDR1^SRFR1^ condensation, we asked whether P1 or P3 may have long range electrostatic interactions with the positively charged P5 segment. Interestingly, P5 mutations K260E/K263E reversed the effect of P1 mutations E148K/E152K, exhibiting dispersed localization (fig. S9B). In contrast, P5 mutations K260E/K263E failed to reverse the effect of P3 mutations D235H/D238H. Because the acidic amino acids in the P1 core sequence SEKSDEKSDK are interspersed by serine and lysine, the long-range electrostatic interaction between P1 and P5 suggests a transient helix formation for P1 segment, with its negatively charged amino acids arranged on one side (fig. S9A). In an unstructured state, the positively charged lysine residues in P1 would neutralize their neighboring negatively charged amino acids. Such transient and metastable interactions in IDRs are well-positioned for fine-tuning protein functions and open up possibilities for induction of helices through protein interactions or post-translational modifications. Combinatorial mutations in P1 and P3 resulted in increased condensate formation (fig. S9B), suggesting that P1 and P3 may function additively or cooperatively in modulating PANT polymerization.

Based on the above findings, we generated P1 and P3 quadruple mutations in the 9 kb *SRFR1* genomic clone *YFP-HA-gSRFR1* and investigated their effects on primary root growth and biomolecular condensate formation in LRCs after transformation of *srfr1-1*. For simplicity, transgenic lines with P1 (E148K/E152K/D155N/D158N) and P3 (D235H/D238H/E242H/D245K) quadruple mutant constructs are designated as *p1* and *p3*, respectively. Surprisingly, primary roots of all three *p1* transgenic lines were significantly longer than those of WT lines (Fig. 4B and fig. S9C). The protein abundance of *p1* mutant line #7 and line #11 was comparable to or slightly lower than that of WT line #7 (fig. S9C), even though the primary root length of *p1* mutant line #7 and line #11 increased by 23% and 17%, respectively, compared with WT line #7. Protein abundance in *p1* line #10 was slightly higher than in WT #7 and its primary root length increased by 45%. Consistently, more LRC condensates were observed in *p1* line #10 (fig. S9D). All three tested *p3* mutant lines showed WT root lengths even though their protein abundances and number of LRC condensates were significantly lower (Fig. 4C, fig. S9, C and D), suggesting that formation of larger aggregation-like puncta, most likely indicating reduced condensate fluidity, can compensate for the lower protein abundance caused by P3 mutation. Further, we measured condensate fluidity by fluorescence recovery after photo- bleaching (FRAP). Signal recovery was detected as early as 30s post-bleaching for condensates formed by wild-type SRFR1 (fig. S9E). In contrast, hardly any fluorescence recovery was observed for most *p1* mutant condensates at 2 minutes post-bleaching (fig. S9E), suggesting lower fluidity. SRFR1 condensates were sensitive to growth conditions and hormone treatment (Fig. 1E, fig. S4 and fig. S10A). We next asked whether condensates formed by SRFR1 variants were still responsive to stimuli. Interestingly, condensates formed by all SRFR1 variants were eliminated by ACC treatment irrespective of differences in fluidity (fig. S10B), suggesting that SRFR1 condensate is also tightly regulated extrinsically, possibly by chaperones and protein degradation. Taken together, these results suggest a link between primary root length and LRC SRFR1 condensate properties, including number, size, and fluidity, and provide a fine-tuning strategy to engineer plants with longer roots by modifying LRC SRFR1 condensate properties.

### Interaction between a polymerization domain and its adjacent IDR1 regulates biomolecular condensate formation

Next, we used purified proteins to test our hypothesis that condensate formation occurs when IDR1^SRFR1^ curtails PANT polymerization. Consistent with *in vivo* data (Fig. 2B, fig. S5A-D), GST-YFP-PANT formed characteristic fibrils (Fig. 4D, fig. S11, A and B, and movie. S3) when treated with 2% PEG even though the protein was fused to a large and soluble GST-YFP tag.

These data also suggest that PANT fibril formation is spontaneous and does not require additional energy input. Under these conditions (PBS buffer + 2% PEG), GST-YFP-PANT-IDR remained soluble; condensates were only observed after treatment with 3% PEG (fig. S11B). In addition, well-resolved GST-YFP-PANT fibrils and spherical GST-YFP-PANT-IDR1 droplets were reproducibly induced in PBS buffer containing 0.02% Tween-20, which reduces the surface tension of solvent (Fig. 4D). These *in vitro* results extend our conclusion from *in vivo* experiments that the interaction between PANT and IDR1 governs condensate formation in the absence of other cellular proteins.

Because the N-terminal part of IDR1^SRFR1^ (135–213), containing P1 and P2 segments, suppressed PANT condensation in protoplasts as effectively as full-length IDR1^SRFR1^ (fig. S9A), we focused on the interaction between P1 and the PANT domain. We first docked P1 to the PANT dimer with MDockPeP2.(*47*) The top model revealed that P1 binds to a PANT monomer surface not predicted to engage in dimerization or tetramerization (Fig. 4E). As negatively charged amino acids in P1 were found to be crucial in regulating condensate formation in protoplasts (fig. S9A), we mutated arginine 20 residue in the PANT domain potentially mediating the interaction between the PANT and P1 segment. Consistent with this prediction, PANT^R20A^ resulted in puncta in protoplasts (Fig. 4F). Moreover, purified GST-YFP-PANT^R20A^ alone formed spherical droplets *in vitro* (Fig. 4D). These results reinforce the idea that subtle changes in PANT are crucial for condensate formation. Neutralizing the positive charge at R20, either naturally through interaction with P1 or artificially by the R20A mutation, may induce a conformational change that shifts the PANT domain from linear polymerization to condensate formation.

Next, we measured the binding affinity between GST-YFP-His-PANT^WT^ and a synthetic P1 peptide. As expected, the disassociation constant for PANT^WT^-P1 interaction was very high, with an estimated KD = 395 μM (fig. S11C), indicating that binding of P1 to PANT is very weak. The interaction between GST-YFP-His-PANT^R20A^ and P1 peptide was even weaker, with an estimated KD = 956 μM (fig. S11C). Because P1 at high concentrations shifted the pH of our PBST buffer, we were not able to determine the exact KD of the PANT-P1 and PANT^R20A^-P1 interactions. Nonetheless, this result indicates that the P1 segment is a molecular recognition motif (MoRF) involved in interacting with the PANT domain. This notion is also supported by the result that P1 is predicted to be involved in protein binding by five different Critical Assessment of protein Intrinsic Disorder (CAID) predictors (fig. S11D).(*45, 48, 49*) The condensate formation of GST-YFP-PANT^R20A^ prompted us to test whether P1-binding induces GST-YFP-PANT^WT^ to form spherical droplets. Droplet formation by PANT^WT^ was indeed induced by P1 peptide (Fig. 4G). Together with *in vivo* evidence, our *in vitro* condensate assays support the idea that the dynamic interaction between PANT and IDR1^SRFR1^ governs condensate formation. Moreover, these results also indicate that a polymerization capable domain (PD) and its adjacent IDR compose a minimal module for polymerization-mediated biomolecular condensate formation.

Then, we tested whether our minimal module paradigm applies to other proteins known to form condensates mediated by polymerization, such as Dishevelled 2 (DVL2) (*18–20*). DVL2 is composed of three folded domains and three IDRs with the arrangement DIX-IDR1-PDZ-IDR2- DEP-IDR3 (fig. S12A). The DIX domain is known to mediate polymerization (*50*). Thus, we assessed the ability of DIX-IDR1^DVL2^ to form condensates and of dehydrins to replace IDR1^DVL2^ (fig. S12, B, C, and S13A). Similarly, segments modulating DIX-IDR1^DVL2^ condensate properties were also identified in IDR1^DVL2^ (fig. S12, D and E). We reasoned that mutations in IDR^DVL2^ should rescue condensate formation of DIX variants if the interaction between DIX and IDR1^DVL2^ governs condensate formation. For example, Y27S and K68T mutations in DIX abolished the formation of long fibrils (fig. S13, B and C). Indeed, fusion of IDR1^DVL2-WT^ to these DIX variants exacerbated the lack of condensates, with DIX^Y27S^-IDR1^DVL2-WT^ and DIX^K68T^-IDR1^DVL2-WT^ almost exclusively exhibiting dispersed localization (fig. S12F). Further, mutations or truncations in IDR1^DVL2^ led to recovery of condensate formation driven by DIX^Y27S^ or DIX^K68T^, with punctate localization observed again for DIX^Y27S^-IDR1^DVL2^ ^Del_94-171^, DIX^Y27S^- IDR1^DVL2^ ^Del_94-171^ ^Del_245-263^, and DIX^Y27S^-IDR1^DVL2^ ^P1^ ^mutant^. Similarly, condensate formation was restored for DIX^K68T^-IDR1^DVL2^ ^Del_94-171^ and DIX^K68T^-IDR1^DVL2^ ^P1^ ^mutant^. Interestingly, DIX^K68T^- IDR1^DVL2^ ^Del_94-171^ ^Del_245-263^ showed reversion to linear fibrils. Collectively, these results strongly suggest that the interaction between DIX and IDR1^DVL2^ governs biomolecular condensate formation, in which DIX drives and IDR1^DVL2^ modulates condensate formation. As for IDR1^SRFR1^, these results imply a segment-spacer architecture for IDR1^DVL2^.

## DISCUSSION

Over the past several decades, considerable efforts have been directed towards identifying genetic mechanisms responsible for plant survival and growth promotion under severe stresses (*51, 52*). However, less attention has been paid to understanding how plants buffer milder environmental fluctuations and genetic variations that nevertheless impact plant productivity. Also, limited success has been achieved by traditional overexpression or loss-of-function strategies (*51, 52*). Recently, it has been realized that growth inhibition is not due to limits in energy supply but is actively initiated by stress signaling pathways, i.e. stress responses are composed of both canonical stress responses and active growth inhibition (*51–53*). To maximize survival, plant genetic programs tend to overcompensate under mild stresses (*51, 52*). Given that extreme or ideal conditions are rare, fluctuating mild stresses, such as daily fluctuations of temperature, humidity, and light intensity, represent significant factors constraining plant growth. Thus, rebalancing the growth-stress tradeoff by avoiding excessive growth inhibition under mild stresses while maintaining a robust stress response under severe conditions should be an effective strategy for improving plant resilience.

In this study, we demonstrated that SRFR1 condensates are required to effectively fine-tune stress responses for optimal growth (Fig. 1E and fig. S4). We showed that *in planta* SRFR1 condensates dynamically respond to varying plant growth conditions (Fig. 1E and fig. S4). Under favorable conditions, large numbers of SRFR1 condensates facilitate rapid root growth.

Conversely, under moderate stress conditions, SRFR1 condensates are reduced to achieve a moderate stress response. When facing extreme conditions (mimicked by ACC treatment), SRFR1 condensates are eliminated to orchestrate a robust stress response. The condensate- independent function of SRFR1 in ETI and the condensate-dependent function of SRFR1 in root development enabled us to finetune its condensate properties to obtain long roots without causing any side effects. Indeed, we have found that the primary root length can be further increased when the fluidity of SRFR1 condensates is reduced by mutating 4 acidic amino acids in the P1 segment of IDR1^SRFR1^ (Fig. 4B and fig. S9C). Moreover, unlike pathological fibrils or aggregations found in human diseases, SRFR1 condensates with reduced fluidity nevertheless can be completely eliminated when necessary (fig. S10B). Based on our results, we consider it likely that a less fluid SRFR1 condensate enables plants to avoid excessive growth inhibition under mild stresses (e.g. horizontal growth) while still effectively responding to severe stresses (e.g. mimicked by ACC treatment).

There is great need for experimentally generalizable toolkits for decoding ensembles of IDRs/IDPs. Over 40% of eukaryotic proteins contain at least one IDR of 30 residues or longer (*45, 54*). Remarkably, approximately 20% of human diseases can be directly linked to mutations occurring in IDRs (*55*). Despite the significance of IDRs in disease, ensembles and functions of IDRs are not well characterized experimentally due to technical challenges. Available bioinformatic and AI tools, including AlphaFold2 and ESM-2, performed well for folded domains (*39, 56*). However, they provide insufficient and inaccurate predictions on IDRs. In addition, biophysical characterization of IDRs/IDPs is time consuming and technically challenging.

Our study on IDR1^SRFR1^ also provides an alternative strategy for decoding ensembles and functions of IDRs. According to the sequence-ensemble-function paradigm (*57–59*), we assume that two IDRs have similar ensembles if they are functionally exchangeable. Thus, if an IDR with unknown function is functionally substituted by a well-characterized IDR, we can examine possible features of the unknown IDR in relation to the characterized IDR. In this study, using protoplast-based high throughput localization screening, we found that type II LEA proteins, but not other types of LEA proteins and prion-like IDRs, closely mimicked the function of IDR1^SRFR1^ (Fig. 3B). These findings led us to examine potential segments that form transient secondary structures in IDR1^SRFR1^. Together with a peptide docking program, *in vitro*, and *in vivo* experiments (Fig. 4E-G and fig. S9A), we were able to identify the P1^SRFR1^ segment as a MoRF. Therefore, this study provides a proof-of-concept for building a toolkit containing multiple IDRs with known functions and ensemble properties to decode unknown IDRS/IDPs.

Together with bioinformatic tools and a high-throughput screening strategy, we believe this experimental toolkit will greatly promote the characterization of IDRs with unknown functions and features.

A key role for the LRC in controlling meristem size has been established by several studies (*38, 60–63*), and the root cap not only protects the meristem from mechanical damage and pathogens, but also is the site for perception and integration of diverse external and internal stimuli (*38, 60, 62, 63*). We hypothesize that control of root growth by LRC SRFR1 condensates may include key regulators or indirectly the translation of key components from multiple signaling pathways. A future unbiased proteomic and translatomic approach is needed to address the mechanistic question of how LRC SRFR1 condensates regulate root growth.

## Acknowledgments

We are grateful to Dr. Alexander Jurkevich and Dr. Frank Baker in the University of Missouri Advanced Light Microscopy Core for their help with confocal microscopy. We thank ABRC for providing Arabidopsis *etr1-3* and *ein2* mutants and plasmids. We thank Longfei Wang from the University of Missouri for their assistance with using SAS for statistical analysis. We also thank Sharon Pike for her comments and editing of the manuscript. This work is supported by National Science Foundation grant IOS-1456181 (WG), USDA-NIFA grant PBI 2022-11933 (WG), National Institutes of Health grant R35GM136409 (XZ) and National Institutes of Health grant 1R01HL119053 (SYC).

## Author contributions

J.S. and W.G. conceived the project, designed experiments, and wrote the manuscript. J.S. performed plasmid construction, genetic transformation, protoplast transfection, confocal microscopy, RT-qPCR, and root measurement. X.X. and X.Z. performed protein structure prediction and molecular docking. L.C. generated protein blots. S.W., Z.D. and W.P. helped with protein purification. R.Z. and S.C. performed HEK293T cell transfection. Z.Z. and B.Y. helped with protoplast preparation and genotyping. K.S. helped with protein-peptide binding analysis.

## Data and materials availability

All data are available in the main text or the supplementary materials.

## Materials and Methods

### *Arabidopsis thaliana* and growth conditions

*Arabidopsis thaliana* ecotype accessions Col-0, RLD, mutants *srfr1-1*, *srfr1-2*, *srfr1-4*, *snc1-11*, *eds1-2*, *srfr1-4 snc1-11* and *srfr1-4 eds1-2* were described previously (*33–36*). *etr1-3* (CS3070) and *ein2* (CS3071) were obtained from Arabidopsis Biological Research Center (ABRC). SRFR1 CRISPR mutants in Col-0 (*srfr1^Col-0^*), RLD (*srfr1^RLD^*) and Ws (*srfr1^Ws^*) background were generated in this study, desired mutations were confirmed by Sanger sequencing. *srfr1-4 etr1-3* and *srfr1-4 ein2* were generated by standard genetic crossing; double mutants were characterized by PCR-based genotyping and conformed by Sanger sequencing. YFP-HA-gSRFR1^WT^; *srfr1-1*, YFP-HA-gSRFR1^Del_PANT^; *srfr1-1*, YFP-HA-gSRFR1^Del_PANT-IDR1^; *srfr1-1*, YFP-HA-gSRFR1^Del_IDR2^; *srfr1-1*, YFP-HA- gSRFR1^WT^; *srfr1-4*, YFP-HA-gSRFR1^Del_PANT^; *srfr1-4*, YFP-HA-gSRFR1^Del_PANT-IDR1^; *srfr1-4*, YFP-HA- gSRFR1^Del_IDR2^; *srfr1-4*, YFP-HA-gSRFR1^WT^; *srfr1^Ws^*, YFP-HA-gSRFR1^Del_PANT^; *srfr1^Ws^*, YFP-HA- gSRFR1^Del_PANT-IDR1^; *srfr1^Ws^*, YFP-HA-gSRFR1^L13T^ ^F47T^ ^K101A^; *srfr1-1*, YFP-HA-gSRFR1^F47T^ ^E115A^; *srfr1-1*, YFP-HA- gSRFR1^IDR1-HIRD11^; *srfr1-1*, YFP-HA-gSRFR1^IDR1-ERD14^; *srfr1-1*, YFP-HA-gSRFR1^IDR1-COR47^; *srfr1-1*, YFP-HA- gSRFR1*^p1^* ^(E148K^ ^E152K^ ^D155N^ ^D158N)^; *srfr1-1*, YFP-HA-gSRFR1*^p3^* ^(D235H^ ^D238H^ ^E242H^ ^D245K)^; *srfr1-1*, were generated in this study (see below for more details).

For root length measurement, if not specified, *Arabidopsis thaliana* seeds were germinated and grown on half-strength Murashige and Skoog (1/2 MS) medium containing 1% (w/v) sucrose, 0.05% (w/v) 2-(N-morpholino) ethanesulfonic acid (MES) and 0.5% agar (pH = 5.7) in transparent petri-dishes placed horizontally on a rack under constant light (75 μE m^-2^ s^-1^) at 23 °C. For the vertical growth assay, *Arabidopsis thaliana* seeds were germinated and grown on 1/2 MS medium containing 1% (w/v) sucrose, 0.05% MES and 1.2 % agar (pH = 5.7). For liquid growth assays, *Arabidopsis thaliana* seeds were germinated and floated on 1/2 MS medium containing 0.25 % (w/v) sucrose and 0.05% (w/v) MES (pH = 5.7). Seedlings from day 6 to day 8 were used for root length measurement and confocal imaging. For the RT-qPCR experiment, seedlings grown in liquid medium were transferred to fresh liquid medium on day 6, root and shoot tissues were harvested on day 12. For transgenic plant screening, *Arabidopsis thaliana* seeds were germinated and grown on half-strength 1/2 MS medium plates containing 1% (w/v) sucrose, 0.05% (w/v) 2-(N-morpholino) ethanesulfonic acid (MES), 0.5% agar (pH = 5.7), 25 μg/ml kanamycin and 100 μg/ml timentin. For seed propagation, all *Arabidopsis thaliana* plants were grown in soil (Beger BM2 seed germination mix) in a Conviron growth chamber at 23 °C, 50% relative humidity, and 120 μE m^-2^ s^-1^ light with a 16h/8h light/dark photoperiod. For protoplast preparation, seeds of Arabidopsis were directly sown on soil and grown in a Conviron reach-in growth chamber at 22 °C, 75% relative humidity, and 100 μE m^-2^ s^-1^ light with a 12h/12h light/dark photoperiod.

### Rice and growth conditions

Rice seedlings were grown on 1/2 MS medium containing 1% (w/v) sucrose and 0.8% agar in a transparent plastic cup under constant light (200 μE m^-2^ s^-1^) at 30 °C. 7-10-day-old seedlings were used for protoplast isolation.

### Bacterial strains

*Agrobacterium tumefaciens* GV3101 harboring desired plasmids was grown on Luria-Bertani (LB) medium (10 g yeast extract, 5 g NaCl, 10 g tryptone, and 12 g agar for 1 L) plates with 25 μg/ml gentamicin and 50 μg/ml kanamycin at 30 °C. *Escherichia coli* BL21 (DE3) pLysS strain containing desired plasmids was grown in LB liquid medium with 34 μg/ml chloramphenicol and 100 μg/ml carbenicillin at 37 °C until an optical density (OD) at 600 nm of 0.4 was reached. Then, cell cultures were incubated on ice for 1 hour before adding 200 mM IPTG. Protein induction was performed by growing cell cultures at 22 °C for 16 hours.

### Human HEK293T cell line

HEK-293T was obtained from American Tissue Culture Collection (ATCC, CRL-3216 ™) and cultured in Dulbecco’s modified Eagle’s medium (DMEM) supplemented with 10% fetal bovine serum (FBS), penicillin (100 U/ml) and streptomycin (100 μg/ml). HEK-293T was transfected with JetPRIME® transfection reagent (101000046) according to the manufacturer’s protocol. Transfected cells were incubated at 37 °C for 36 hours before confocal imaging.

### Plasmid construction and transgenic plant preparation

*SRFR1* CRISPR construct was generated according to our previous work (*64*). Briefly, two guide RNAs targeting *SRFR1* exon 3 and exon 6 and driven by Arabidopsis U6-26 and U3b promoters were cloned to *pAtEC1.2e1p::Cas9-GFP-gRNA* vector. *SRFR1* CRISPR mutants in Col-0, RLD and Ws background were generated by *Agrobacterium tumefaciens* GV3101-mediated transformation. Positive transformants were selected by spraying Basta and confirmed by Sanger sequencing. Eventually, the same 1 bp insertion allele in all backgrounds was used for further analysis.

For generating YFP-HA tagged SRFR1 transgenic lines, a previously reported HA-tagged 9 kb SRFR1 genomic fragment in *pCambia2300-HA-gSRFR1* was modified by overlap PCR to introduce AatII and SpeI cloning sites; then the YFP tag was inserted between these two sites just before the HA tag. This construct was designated as *pCambia2300-YFP-HA-gSRFR1^WT^*. Subsequently, SRFR1 domain truncation binary constructs were generated by overlap PCR. To make the PANT domain truncation construct (SRFR1^Del_6-134^), a ∼930 bp fragment 1 was PCR amplified with PANTdel-LP + PANTdel-Overlap-RP. A 756 bp fragment 2 was obtained with PANTdel-Overlap- RP + PANTdel-RP. Using fragment 1 and 2 as template, PCR with PANTdel-LP + PANTdel-RP was used to amplify a ∼1660 bp fragment. This band was cut with AatII and XbaI to recover an ∼1200 bp band and ligate it to AatII-and XbaI-cut *pCambia2300-YFP-HA-gSRFR1^WT^*. Then we used PANTdel-RP to sequence the construct.

Correspondingly, the PANT-IDR1 deletion (SRFR1^Del_6-289^) construct was generated with primer combinations of PANTdel-LP + PANT-IDR1del-Overlap-RP, PANT-IDR1del-Overlap-LP + PANT-IDR1del-RP, and PANTdel-LP + PANT-IDR1del-RP. The resulting fragment was cut with AatII and XbaI. A ∼300 bp band was recovered and ligated to AatII- and XbaI-cut *pCambia2300-YFP-HA-gSRFR1^WT^*. IDR2 deletion (SRFR1^Del_715-751^) was generated with primer combinations of IDR2del-LP + IDR2del-Overlap-RP, IDR2del-Overlap-LP + IDR2-del-RP, and IDR2del-LP + IDR2-del-RP. The resulting fragment was digested with KfII and BstBI; an ∼1800 bp band was recovered and ligated to KflI- and BstBI-cut *pCambia2300-YFP-HA-gSRFR1^WT^*. All above constructs were sequenced and transformed to *srfr1-1* and *srfr1-4* via *Agrobacterium tumefaciens* GV3101-mediated transformation. T3 homozygous plants were characterized and used for further analysis.

To generate *pCambia2300-YFP-HA-gSRFR1^L13T^ ^F47T^ ^K101A^* and *pCambia2300-YFP-HA-gSRFR1^F47T^ ^E115A^* constructs, an ∼2 kb fragment covering YFP-HA and SRFR1 exons 1-3 was subcloned to pBlueScript II SK (+) and designated as *pBS-YFP-HA-SRFR1^Exon1-3^*. PCR-based site-specific mutagenesis was performed to make *pBS-YFP-HA-SRFR1^Exon1-3^ ^L13T^ ^F47T^ ^K101A^* and *pBS-YFP-HA-SRFR1^Exon1-3^ ^F47T^ ^E115A^*. After sequencing, ∼2 kb fragments harboring desired mutations were cut out with AatII and XbaI and ligated to AatII- and XbaI-digested *pCambia2300-YFP-HA- gSRFR1^WT^*. To make *pCambia2300-YFP-HA-gSRFR1^IDR1-HIRD1^*, *pCambia2300-YFP-HA-gSRFR1^IDR1-ERD14^*, *pCambia2300-YFP-HA-gSRFR1^IDR1-COR47^*, *pCambia2300-YFP-HA-gSRFR ^p1^ ^(E148K^ ^E152K^ ^D155N^ ^D158N)^*, and *pCambia2300-YFP-HA-gSRFR1^p3^ ^(D235H^ ^D238H^ ^E242H^ ^D245K)^* constructs, MluI and XmaI sites were first introduced to *pBS-YFP-HA-SRFR1^Exon1-3^* through overlap PCR to get *pBS-YFP-HA-SRFR1^Exon1-3^ ^MluI-XmaI^*. Then, coding sequences (without stop codon) of HIRD11, ERD14, COR47, IDR1^SRFR^ *^p1^* ^(E148K^ ^E152K^ ^D155N^ ^D158N)^ and IDR1^SRFR1^ *^p3^* ^(D235H^ ^D238H E242H D245K)^ were cut with MluI and XmaI from their *pSAT6-eYFP* constructs and ligated to MluI- and XmaI-digested *pBS-YFP-HA-SRFR1^Exon1-3^ ^MluI-XmaI^*. After sequencing, ∼2 kb fragments harboring desired mutations were cut out with AatII and XbaI and ligated to AatII- and XbaI-digested *pCambia2300-YFP-HA-gSRFR1^WT^*.

To generate *pSAT6* based transient expression SRFR1 and DVL2 truncation constructs, fragments corresponding to full-length SRFR1, PANT, IDR1^SRFR1^, PANT-IDR1^SRFR1^, IDR1^SRFR1^-SMiT, SMiT, SRFR1^N^ (PANT- IDR1-SMiT), SRFR1^C^ (SACT1-IDR2-SACT2), SRFR1^Del_PANT^ (IDR1-SMiT-SACT1-IDR2-SACT2), full-length DVL2, DIX, IDR1^DVL2^, and DIX-IDR1^DVL2^ were amplified with PCR (for primer details, see Table S1), digested with KpnI and XmaI, and ligated to KpnI- and XmaI-cut *pSAT6-eYFP* (ABRC: CD3-1103) and *pSAT6-mRFP* (ABRC: CD3-1107) as indicated. cDNA fragments for SRFR1^Del_IDR1^ and SRFR1^Del_IDR2^ were generated by overlap PCR. The ∼1890 bp cDNA SRFR1^Del_IDR1^ (SRFR1^Del_142-294^) fragment was obtained using primer combinations of IDR1del-CDS-LP + IDR1del-CDS-Overlap-RP, IDR1del-CDS-Overlap-LP + IDR1del-CDS-RP, and of IDR1del- CDS-LP + IDR1del-CDS-RP. The fragment was then cut with KpnI and AflII. A 1552 bp fragment was recovered and ligated to KpnI- and AflII-cut *pSAT-eYFP-SRFR1*. ∼1600 bp cDNA SRFR1^Del_IDR2^ (SRFR1^Del_718-754^) fragment was obtained using primer combinations of IDR2del-CDS-LP + IDR2del-CDS-Overlap-RP, IDR2del-CDS-Overlap- LP + IDR2del-CDS-RP, and IDR2del-CDS-LP + IDR2del-CDS-RP. After this fragment was cut with AflII and NotI, an ∼1380 bp fragment was recovered and ligated to AflII- and NotI-cut *pSAT-eYFP-SRFR1*. Stop codon was incorporated into all fragments if not specified, i.e., no additional sequence was added due to cloning.To generate PANT and IDR fusion constructs, a short fragment containing MluI and XmaI was introduced to *pSAT6-eYFP-PANT-Stop* to get *pSAT6-eYFP-PANT-No Stop (NS)*. A 10 amino acid tail was added after the PANT domain due to the multiple cloning site in *pSAT6-eYFP-PANT-NS*. Similarly, a short fragment containing MluI and XmaI was introduced to *pSAT6-eYFP-DIX-Stop* to get *pSAT6-eYFP-DIX-No Stop (NS)*. Coding sequences corresponding to IDR1^NtSRFR1^, IDR1^GmSRFR1^, IDR1^ZmSRFR1^, IDR1^OsSRFR1^, IDR1^AtSRFR1^, IDR1^SRFR1 Del_135-212^, IDR1^SRFR1 Del_214-289^, IDR1^SRFR1 Del_135-212 Del_273-289^, IDR1^DVL2 Del_94-171^, IDR1^DVL2 Del_245-263^, IDR1^DVL2 Del_94-171 Del_245-263^, HIRD11, XERO1, XERO2, COR47, ERD14, COR15A, LEA4-1, IDRFLL2, IDRHEM1, IDRFCA, and IDR1TDP43 were obtained by PCR. These fragments then were cut with MluI and XmaI and ligated to MluI- and XmaI-cut *pSAT6-eYFP-PANT-NS* and *pSAT6-eYFP-DIX-NS* as indicated. All constructs were confirmed by Sanger sequencing. PANT-HIRD11 fusion with HIRD11 truncation variants were generated similarly. Point mutation variants of PANT, PANT-IDR1, DIX, and DIX-IDR1 were generated by PCR-based site-specific mutagenesis in *pSAT6-eYFP-PANT-Stop*, *pSAT6-eYFP- PANT-IDR1^SRFR1^-NS*, *pSAT6-eYFP-DIX-Stop*, *pSAT6-eYFP-DIX-IDR1^DVL2^-Stop* with primers listed in Table S1. All constructs were confirmed by Sanger sequencing.

To generate pGEX4T-3-YFP, pGEX4T-3-YFP-PANT, pGEX4T-3-YFP-PANT-IDR1^SRFR1^ ^135-289^, fragments of YFP, YFP-PANT, YFP-PANT-IDR1^SRFR1^ ^135-289^ were obtained by PCR using pSAT6-eYFP, pSAT6-eYFP-PANT-Stop, pSAT6-eYFP-PANT-IDR1-NS as templates, then digested by EagI and ligated to SmaI- and EagI-cut pGEX4T-3. pGEX4T-3-YFP-6His-PANT constructs were generated by inserting a short fragment encoding 6His between BglII and SalI in pGEX4T-3-YFP-PANT. pGEX4T-3-YFP-6His-PANT^R20A^ was generated by PCR-based site-specific mutagenesis using pGEX4T-3-YFP-6His-PANT as template. To generate pCMV-based constructs, fragments of YFP-tagged full-length SRFR1, PANT, PANT-IDR1^SRFR1^, IDR1^SRFR1^, full-length DVL2, DIX, DIX-IDR1^DVL2^ and IDR1^DVL2^ were amplified by PCR (for primer details, see Table S1). Then, these fragments were digested with AsiSI and SmaI and ligated to AsiSI- and PmeI-cut pCMV plasmid. All constructs were confirmed by Sanger sequencing.

### Protoplast isolation and transfection

Arabidopsis protoplast isolation and plasmid transfection were strictly conducted according to a previous protocol (*65*). Fully expanded leaves from 5-week-old soil grown Col-0 were used for protoplast isolation. Leaf strips were digested with enzyme solution (1.5% Cellulase R-10, 0.4% Macerozyme R-10, 0.4 M mannitol, 20 mM KCl, 20 mM MES at pH 5.7, 10 mM CaCl2 and 0.1% BSA) in dark for three hours at room temperature. Then, the enzyme solution was diluted with an equal volume of W5 solution (154 mM NaCl, 125 mM CaCl2, 5 mM KCl and 2 mM MES at pH 5.7). Protoplasts were released by gentle shaking. After washing with W5 solution, protoplasts were resuspended in MMG solution (0.4 M mannitol, 15 mM MgCl2 and 4 mM MES at pH 5.7) at a concentration of 2 × 10^5^ protoplasts per ml. 10 μg plasmid DNA per 100 μl protoplast suspension was mixed followed by adding 110 μl PEG solution (40% PEG4000, 0.2 M mannitol, and 100 mM CaCl2). After 15 min incubation, transfection was stopped by adding 440 μl W5 solution. Finally, protoplasts were resuspended in 1 ml W5 solution and incubated in 6-well-plates for 12-16 hours before confocal imaging.

Rice protoplast isolation and transfection was modified from our previous protocol (*66*). Rice (*Oryza sativa*) cultivar Kitaake seeds were sterilized with 2.5% sodium hypochlorite containing 0.05% (w/v) Tween 20 for 30 min, washed with sterile water, and then sown on 1/2 MS medium plates and grown under constant light (200 μE m^-2^ s^-1^) at 30 °C for 7-10 days. Rice protoplasts were prepared by incubating stem strips with enzyme solution (1.5% Cellulase RS, 0.75% Macerozyme R-10, 0.6 M mannitol, 10 mM MES at pH 5.7, 10 mM CaCl2 and 0.1% BSA) for 6-8 h in the dark with gentle shaking (40-60 rpm). After the enzymatic digestion, an equal volume of W5 solution (154 mM NaCl, 125 mM CaCl2, 5 mM KCl and 2 mM MES at pH 5.7) was added, followed by vigorous shaking by hand for 10 sec. After washing with W5 solution, protoplasts were resuspended in MMG solution (0.4 M mannitol, 15 mM MgCl2 and 4 mM MES at pH 5.7) at a concentration of 2 × 10^6^ protoplasts per ml. 10 μg plasmid DNA was used for 100 μl protoplasts. Transfection procedures were the same as those for *Arabidopsis protoplasts*. After 12-16 hours incubation, protoplasts were harvested for confocal imaging or Western blot.

### Confocal microscopy

A Leica TCS SP8 laser scanning confocal microscope was used to image protoplasts, live tissues, and *in vitro* fibrils and condensates. For Arabidopsis protoplast imaging, the excitation wavelength for YFP and chlorophyll auto- fluorescence was 514 nm with 10% laser power. 2% laser power was used for fibril imaging. The emission wavelengths for YFP and chlorophyll auto-fluorescence were 525-575 nm and 650-750 nm, respectively. For rice protoplast imaging, the excitation wavelength for YFP was 514 nm with 2% laser power. RFP was excited by a laser line at 588 nm with 10% laser power. The emission wavelengths for YFP and RFP were 525-575 nm and 570-620 nm, respectively. For *in vitro* fibrils and condensates imaging, a YFP laser line at 514 nm with 0.4% laser power was used. Propidium Iodide (PI) staining of Arabidopsis root was performed by immersing whole seedlings in PI staining solution (30 μg/ml PI in liquid ½ MS medium, pH5.7) for 2 min. Then roots were washed in ½ MS medium and subjected to confocal imaging. A 536 nm laser line with 10% laser power was used for PI excitation, and 600- 680 nm wavelengths were used to detect the PI emission signal. Six-day-old seedlings of *YFP-HA-gSRFR1^WT^* line #7 were treated with ACC (1 μM), AVG (1 μM), IBA (1 μM), IAA (1 μM), and BAP (0.1 μM) for 12 hours prior to confocal imaging.

For FRAP analysis, the 488 nm line of the Argon laser with 80% laser power was used for YFP photobleaching. Pinhole was set to 2 AU and the xyt scanning mode was used. PMT detector (510-560 nm) was used to collect emission signals. 5 pre-bleach images every 1 sec was taken at 0.3-0.5% laser transmission. 4 bleaching repeats in a specified ROI with laser transmission set to 100%. Post-bleaching imaging was recorded every 2 sec for 120 sec using the same settings as for pre-bleaching.

### Recombinant protein isolation and binding assay

To ensure high expression of recombinant proteins, single colonies were picked from freshly transformed *E. coli* BL21(DE3) pLysS plates. Around 10 intermediate-sized single colonies were mixed for inoculation and then incubated overnight at 37 °C in LB broth. For GST-YFP-PANT and GST-YFP-6His-PANT purification, 3 L of LB were inoculated with 20 ml of overnight culture and incubated for approximately 3 hours to reach an optical density of 0.4-0.5. This cell culture was then incubated on ice for 1 hour. IPTG, final concentration 200 μM, was added to the cell culture. After growing overnight at 22°C, the cell culture was passed through a French press to lyse the cells. The extract was centrifuged and 3 ml of glutathione agarose resin (GoldBio, cat# G-250) were added to 150 ml supernatant of GST-YFP-PANT. Samples were incubated at 4°C for 1 hour with rotation and then washed with PBS in a gravity column. Protein was eluted from beads with 10 ml of 50 mM reduced glutathione (prepared with PBS, pH adjusted to 7.4). Protein was diluted with PBS, PBST or Tris buffer (Tris-HCl, pH = 7.5, 150 mM NaCl) and then concentrated with Amicon centrifugal filters (10 kDa cut-off) several times until the concentration of glutathione was below 0.5 mM. Purity and the correct size of purified proteins were confirmed by SDS-PAGE and Coomassie blue staining. Because PANT is an aggregation-prone protein when expressed alone, less than 20% is soluble. For this reason, large volumes of cell culture and glutathione agarose resin were used. Still, about 50% GST-YFP-PANT aggregated during the concentration step. Moreover, fine fibrils of GST-YFP-PANT only form with freshly prepared protein. After freezing, even in the presence of 25% glycerol, only aggregation-like structures formed. For purification of GST-YFP, GST-YFP-PANT-IDR1 and GST-YFP-6His-PANT^R20A^, about 1/3 amount of cell culture and glutathione agarose resin were used.

For MST binding analysis, freshly prepared GST-YFP-6His-PANT^WT^ and GST-YFP-6His-PANT^R20A^ were centrifuged for 15 min at 4 °C, 21, 000g to remove spontaneously formed aggregates. Then, both proteins were diluted to 1.5 mM with PBST and incubated with RED-tris-NTA dye at room temperature for 30 min. P1 peptide in a range of concentrations (5 mM to 1.22 μM) was incubated with labelled prepared GST-YFP-6His-PANT^WT^ and GST-YFP-6His-PANT^R20A^ for 30 min at room temperature. Approximately 10 ml of each sample was loaded to a capillary tube. Measurements were performed with Monolith X instrument (Nano Temper Technologies). Ligand (P1 peptide in this study) binding causes a spectral shift of the fluorophore (RED-tris-NTA) that is bound to a target (GST-YFP-6His-PANT^WT^ and GST-YFP-6His-PANT^R20A^). The disassociation constant Kd was derived by plotting the spectral shift of fluorescence signal detected simultaneously at 670 nm and 650 nm against the P1 concentrations (*67*).

### *In vitro* fibril and condensate assay

For the *in vitro* fibril and condensate assay, proteins of GST-YFP, GST-YFP-PANT and GST-PANT-IDR1 were all prepared freshly and diluted with the indicated buffer to 30 μM. Then, these proteins were centrifuged for 15 min at 4 °C, 21, 000g to remove protein aggregates formed during concentration steps. The *in vitro* fibril and condensate assay was performed in a 200 ml PCR tube at room temperature by mixing with 10 × PEG8000 stock solution. To ensure that PEG8000 was completely dissolved, PEG solution freshly prepared with indicated buffers was shaken at 40-60 rpm for at least 1 hour. For example, 40 μl of 30 μM GST-YFP-PANT was added to a PCR tube containing 4 μl of 10% PEG8000 and mixed by gentle tapping. Then, these whole mixtures were transferred to 18 well m-Slides (Ibidi, cat#81816) for confocal imaging. For P1-induced droplet formation, 40 μl of 30 μM GST-YFP-PANT was first incubated with different concentrations of P1 peptide at room temperature for 30 min to reach equilibrium.

Then, these samples were mixed with 4 μl of 10% PEG8000 and incubated at room temperature for 3 hours.

### Total RNA isolation and RT-qPCR analysis

Total RNA from young seedlings of Arabidopsis, rice, soybean, maize, tobacco, and human HEK293T cells was extracted using TRIzol reagent (Invitrogen). Endogenous RNase was further removed by ethanol precipitation. After DNase treatment, 1 μg of total RNA was used for first strand cDNA synthesis with M-MLV reverse transcriptase (Promega) and oligo (dT) 18 primer. Real-time quantitative PCR (RT-qPCR) was performed using SYBR Green qPCR Master Mix (Agilent) in an ABI 7500 real-time qPCR machine (Life Technologies). *EF1a* was used as an internal control.

### Root measurement

Roots of Arabidopsis seedlings were aligned on the surface of Agar plates. Then, images were captured with a digital camera. The length of primary roots was measured using Image J software. To avoid experimental errors, seeds were harvested from plants grown side by side, and different genotypes were grown in the same Petri dish with multiple replicates.

### Protein structure prediction and molecular docking

PANT dimer and tetramer structures were modeled by a locally installed AlphaFold v2.2.4 using default settings (*39*). Interestingly, the predicted tetramer structure contains two modeled PANT dimeric structures, i.e., the modeled PANT dimeric interface was retrieved in the predicted tetramer structure. Therefore, the structure of 24-mer PANT polymer was generated by elongating the tetramer using the MatchMaker function in the UCSF Chimera program (*68*). Peptide structures of P1^SRFR1^-P5^SRFR1^ and P1^DVL2^-P6^DVL2^ in solution were modeled using the PEP-FOLD 3.5 webserver with default settings (*46*). The binding mode of P1^SRFR1^ on the PANT dimer was modeled using the MDockPeP2 webserver with default settings (*47*). The top-ranked structure from each modeling program was presented in this study. Structures of peptides, proteins, and protein-peptide complexes were analyzed and displayed using UCSF Chimera X (*69*).

### Immunoblot

Immunoblot analysis of the expressed HA-tagged SRFR1 variants was performed by extracting approximately 0.15 gram of whole seedlings in 0.3 mL of 2 × sodium dodecyl sulfate (SDS) loading buffer (100 mM Tris–HCl, pH 6.8, 4% w/v SDS, 20% v/v glycerol, 30 mM dithiothreitol). For detection of GFP-tagged SRFR1 and DVL2 variants in rice protoplast, 0.3 ml of 2 × SDS loading buffer was added to 100 ml of protoplast pellet. Samples were cleared by centrifugation at 21,000 × g for 15 min at 4°C and loaded onto an 8% v/v SDS-PAGE gel. Protein was detected with 1:3,000 horseradish peroxidase-conjugated anti-HA (Roche, cat#12013819001) or anti-GFP (Invitrogen, A11122) antibodies.

### Protein accession number

Protein sequences in this study can be found in The Arabidopsis Information Resource (TAIR, www.arabidopsis.org) and UniProt (www.uniprot.org). SRFR1 (AT4G37460), ETR1 (AT1G66340), EIN2 (AT5G03280), SNC1 (AT4G16890), EDS1 (AT3G48090), HIRD11 (AT1G54410), XERO1 (AT3G50980), XERO2 (AT3G50970), ERD14 (AT1G76180), COR47 (AT1G20440), COR15A (AT2G42450), LEA4-1 (AT1G32560), FLL2 (AT1G67170), HEM1 (AT2G35110), FCA (AT4G16280), ACS1 (At3g61510), ACS2 (At1g01480), ACS4 (At2g22810), ACS6 (At4g11280), ACS7 (At4g26200), ACS8 (At4g37770), ACS9 (At3g49700), ACS11 (At4g08040), EF1a (At5g60390), DVL2 (UniProt: O14641), TDP-43 (UniProt: Q13148).

### Statistical analysis

Significant differences were determined using Student’s t-test and one-way ANOVA. Multiple comparisons were conducted using Tukey-Kramer HSD (p < 0.05).

---

**Fig. S1.**
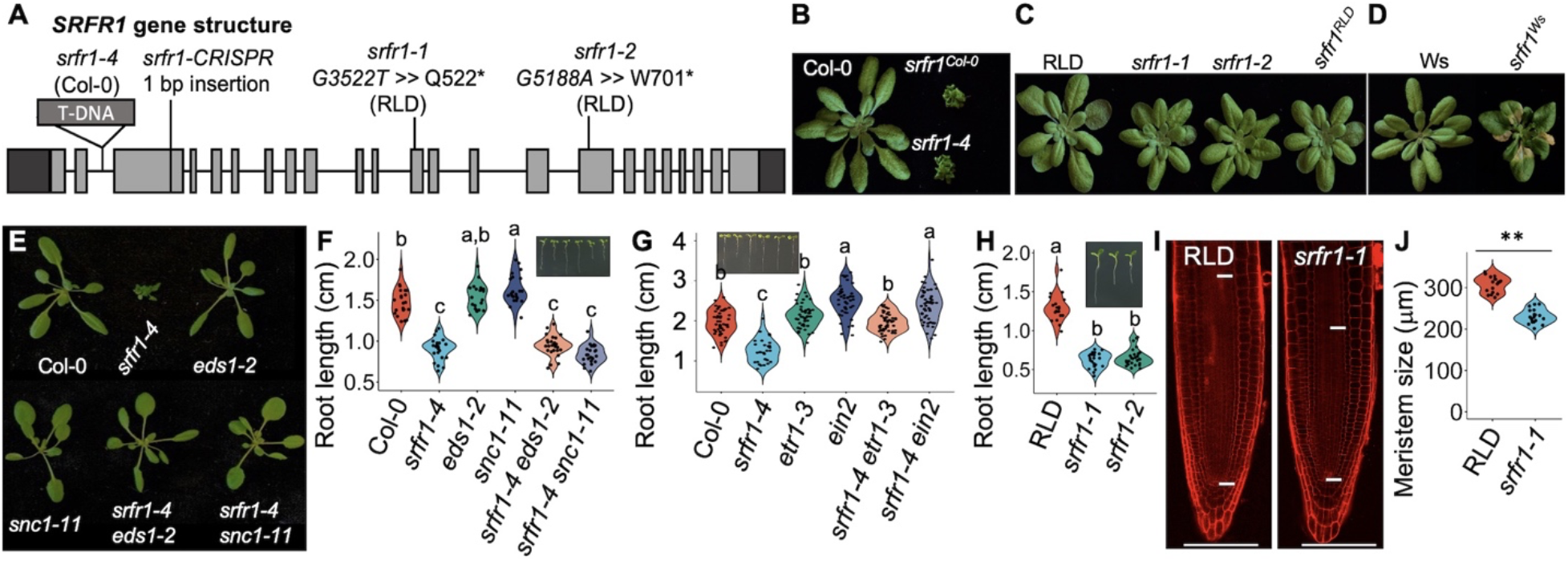
SRFR1 regulates optimal primary root growth. (**A**) Schematic diagram of *SRFR1* gene structure. *srfr1-1* and *srfr1-2* (RLD background) and *srfr1-4* (SAIL_412_E08, Col-0 background) have been described in previous studies (*33, 34*). *SRFR1* CRISPR mutants in Col-0, RLD and WS backgrounds all harbor 1 bp insertion at the same site, designated as *srfr1^Col-0^*, *srfr1^RLD^* and *srfr1^WS^*, respectively. (**B**-**D**) Rosette phenotype of soil-grown *srfr1* mutants in Col-0 (**B**), RLD (**C**), and WS (**D**) backgrounds. (**E** and **F**) The stunting phenotype (**E**), but not the short primary root phenotype (**F**), of *srfr1-4* is dependent on EDS1 or SNC1. Primary root length is measured with 6-day-old seedlings, n = 21-27. (**G**) The short primary root phenotype of *srfr1-4* is dependent on ethylene signaling. Primary root length is measured with 7-day-old seedlings, n = 31-52. (**H**) Primary root length of *srfr1-1* and *srfr1-2*. Primary root length is measured with 6-day-old seedlings, n = 20-28. (**I** and **J**) Root apical meristem size of RLD and *srfr1-1* measured with 6-day-old seedlings, n = 18. ** indicates p < 0.001. Bar = 50 μm.

**Fig. S2.**
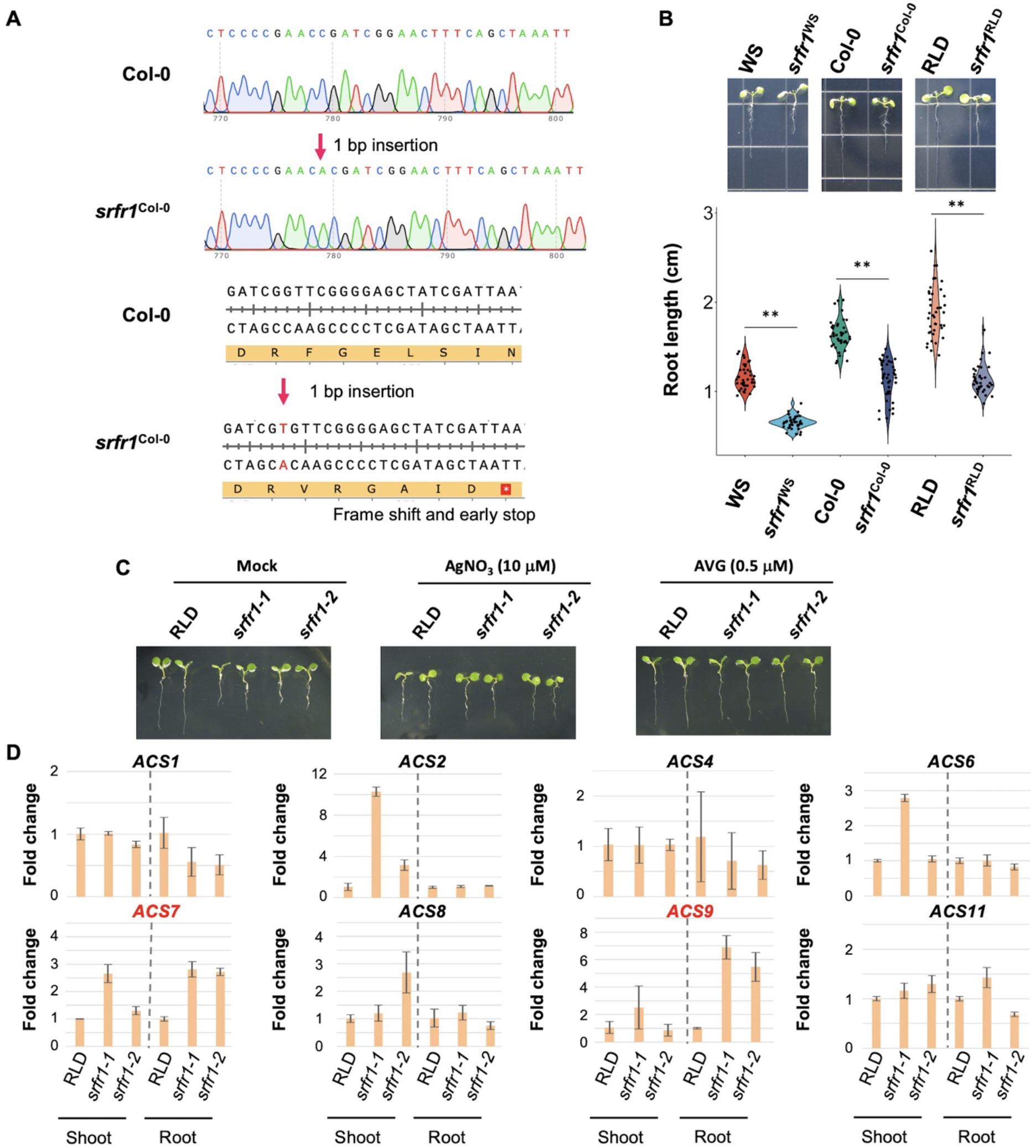
Reduced primary root length in *srfr1* mutants correlates with elevated ethylene response. (**A**) 1 bp insertion in *SRFR1* CRISPR mutants results in a frameshift and an early stop. CRISPR mutants in Col-0, RLD and WS background with the same type of editing were selected for analysis and designated as *srfr1^Col-0^*, *srfr1^RLD^* and *srfr1^WS^*, respectively. (**B**) Primary root length of *SRFR1* CRISPR mutants in Col-0, RLD and WS background. Root length was measured with 6-day-old seedlings, n = 40. Violin-plots show the data distribution. ** indicates p < 0.001. (**C**) The short root phenotype of *srfr1-1* and *srfr1-2* is largely rescued by inhibiting ethylene biosynthesis. Primary root length was measured with 6-day-old seedlings grown on ½ MS plates containing AgNO3 and AVG concentrations as indicated. (**D**) Real-time qPCR analysis of ethylene biosynthesis gene expression in shoot and root tissues of RLD, *srfr1-1*, and *srfr1-2*. Values are means ± SD, *n* = 3. The primers used for real-time qPCR were according to a previous study (*70*). *EF1a* was used as an internal control.

**Fig. S3.**
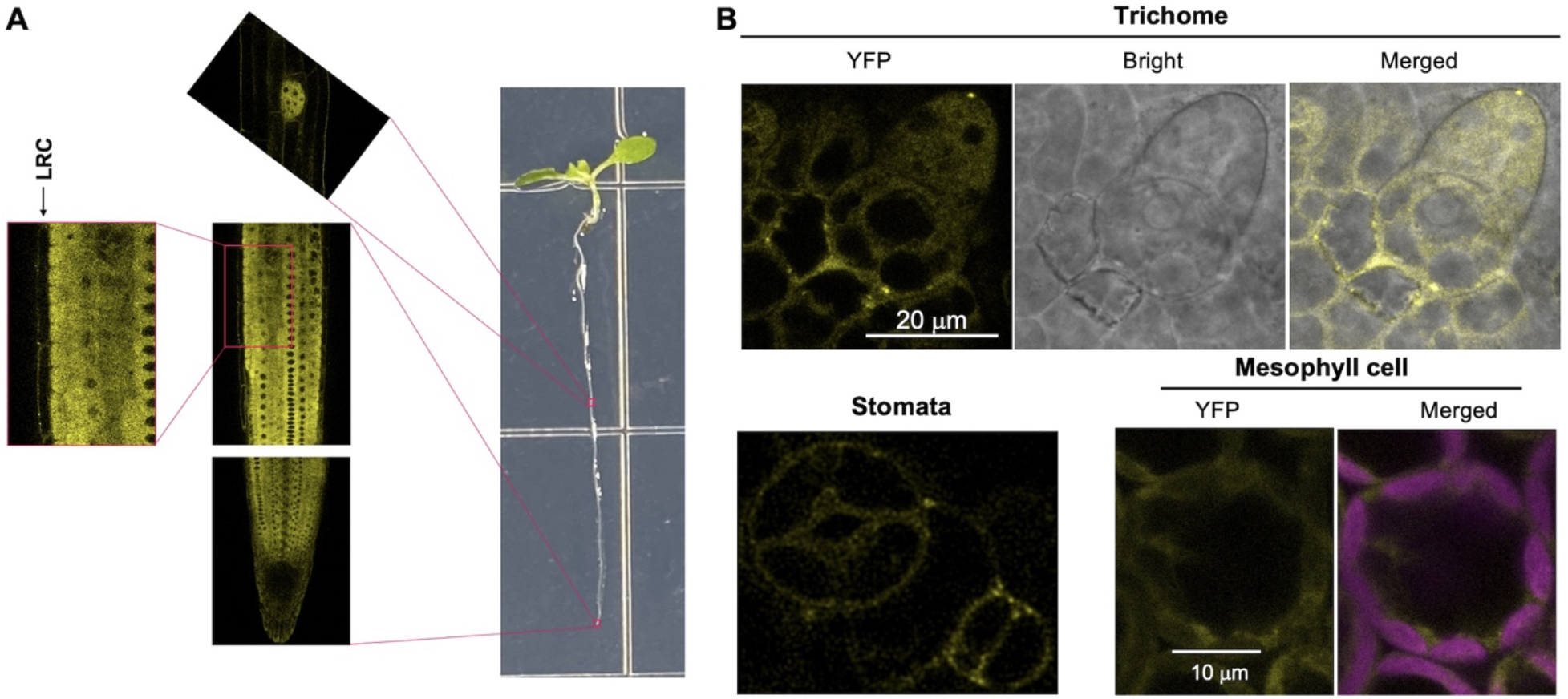
Localization of SRFR1 in stable transgenic plants. (**A**) Subcellular localization of SRFR1 in roots of 6-day-old seedlings. (**B**) Subcellular localization of SRFR1 in true leaf tissues of 8-day-old seedlings.

**Fig. S4.**
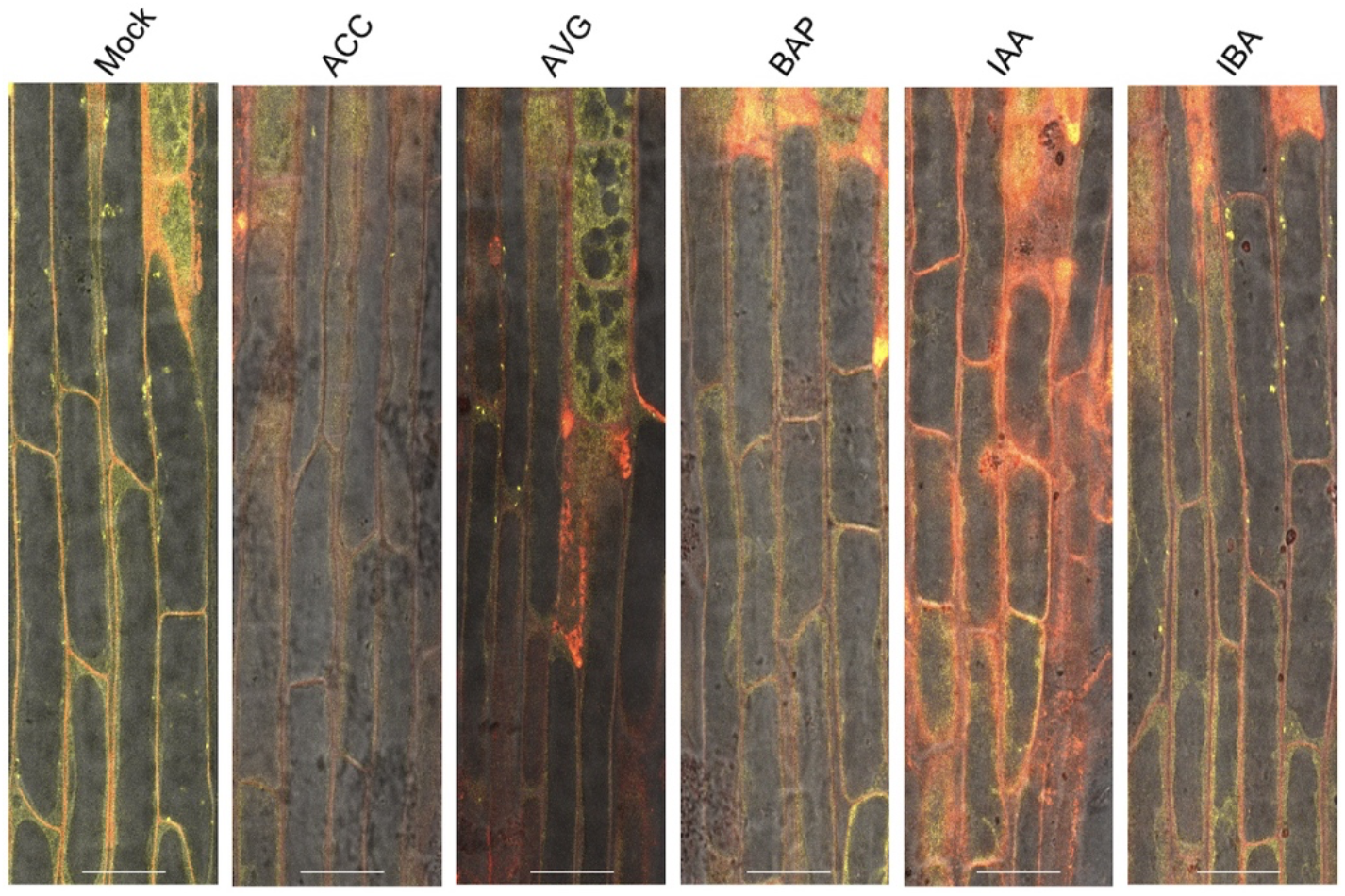
Accumulation of LRC SRFR1 condensates is responsive to hormone treatments. Single median section images were taken with roots of 6-day-old *srfr1-1 YFP-HA-gSRFR1^WT^* #7 seedlings grown under different conditions. (Q) Hormone treatment was performed at day 5 with *srfr1-1 YFP-HA-gSRFR1^WT^* #7 seedlings grown in liquid ½ MS medium. Single median section images were taken at 12 hours post hormone treatment. Bar = 25 μm.

**Fig. S5.**
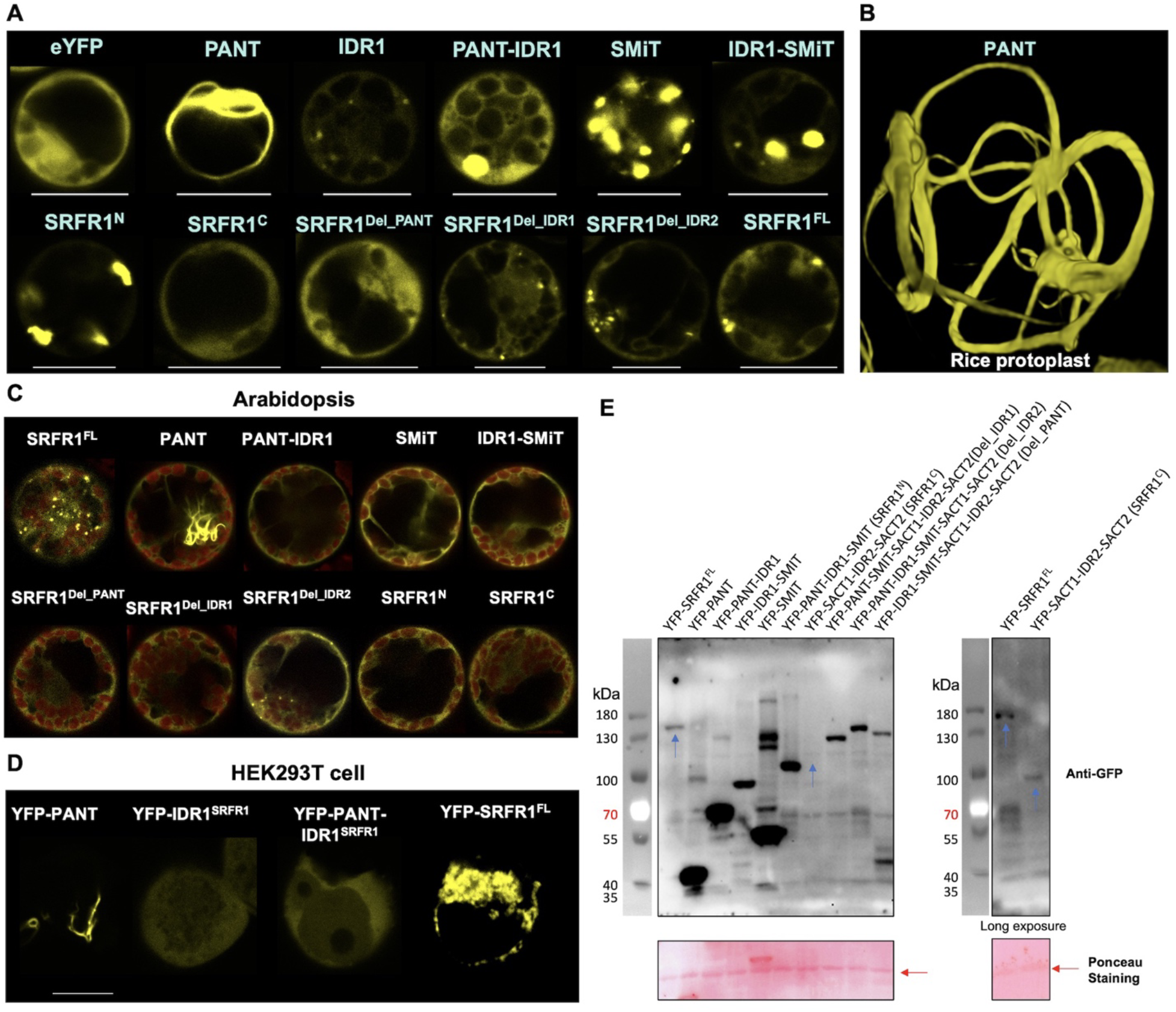
Localization of SRFR1 and its truncated variants in Arabidopsis, rice and human HEK293T cells. (**A** and **B**) Subcellular localization of full-length and differentially truncated SRFR1 in rice protoplasts. Bar = 10 mm. Images were taken with the same confocal microscopy settings. (**C**) The PANT domain is required for SRFR1 biomolecular condensate formation in Arabidopsis protoplasts. SRFR1 and differentially truncated SRFR1 variants were expressed in Arabidopsis protoplasts. (**D**) Subcellular localization of PANT, IDR1^SRFR1^, PANT-IDR1^SRFR1^, and full-length SRFR1 in human HEK293T cells. Bar = 10 mm. (**E**) Western-blot detection of YFP-tagged full-length and truncated SRFR1 expression in rice protoplasts. Ponceau S staining of the Rubisco large subunit is shown as a protein loading control. Blue arrows indicate lower protein accumulation of SRFR1^FL^ and SRFR1^C^.

**Fig. S6.**
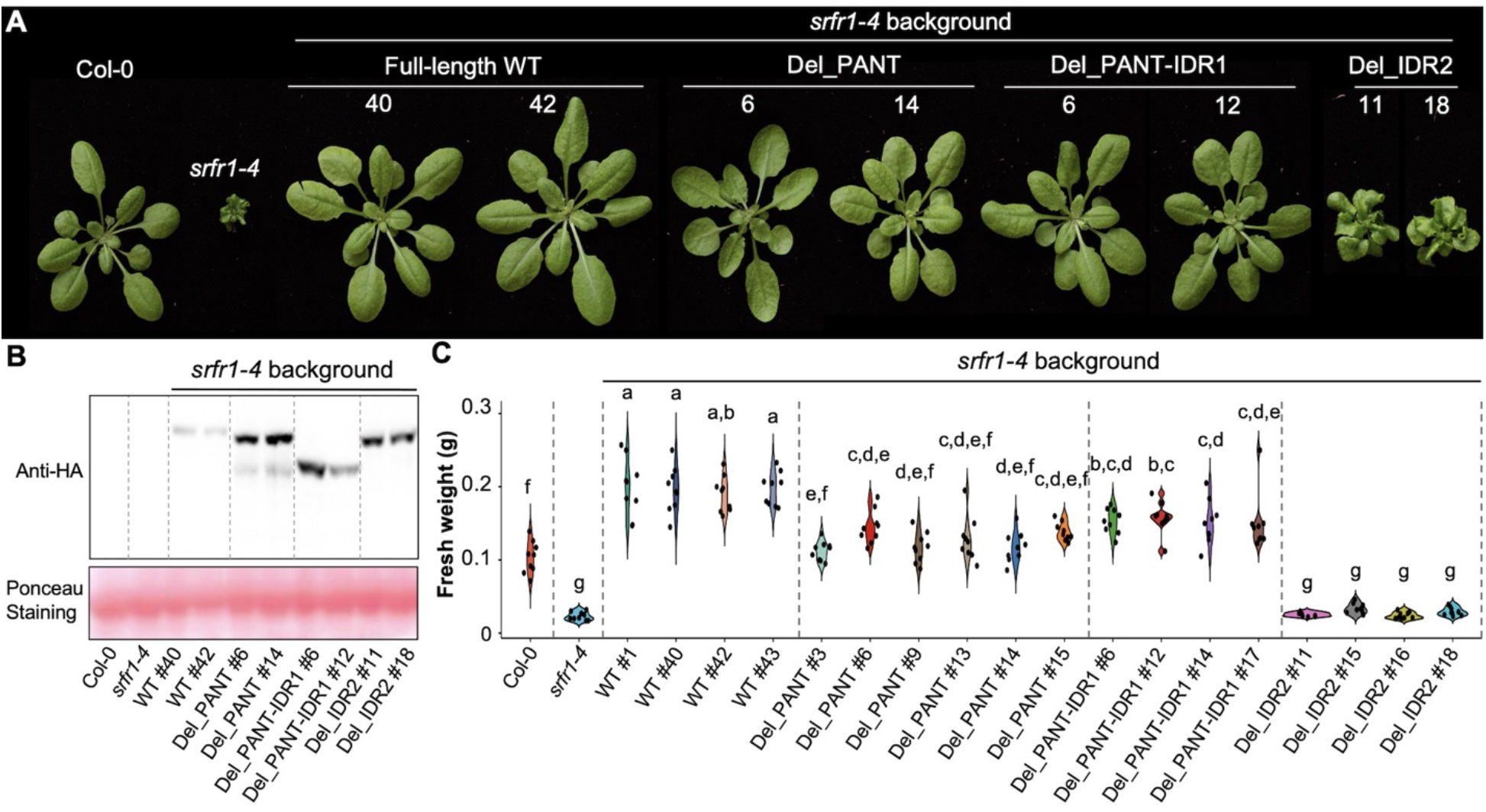
The PANT-IDR1 module is required for shoot growth promotion. (**A**) Rosette phenotype of 4-week-old soil-grown plants. All SRFR1 truncated variants were transformed in the *srfr1-4* background. (**B**) Western-blot detection of YFP-HA-tagged full-length and truncated SRFR1 expression in the *srfr1-4* background. Ponceau S staining of the Rubisco large subunit shows equal protein loading. (**C**) The PANT-IDR1 module is required for increased SRFR1-mediated biomass. Fresh weights of different genotypes were measured with 4-week-old soil-grown plants, n = 9. Violin-plots show data distribution. Letters denote different statistical groups (ANOVA, Tukey-Kramer grouping).

**Fig. S7.**
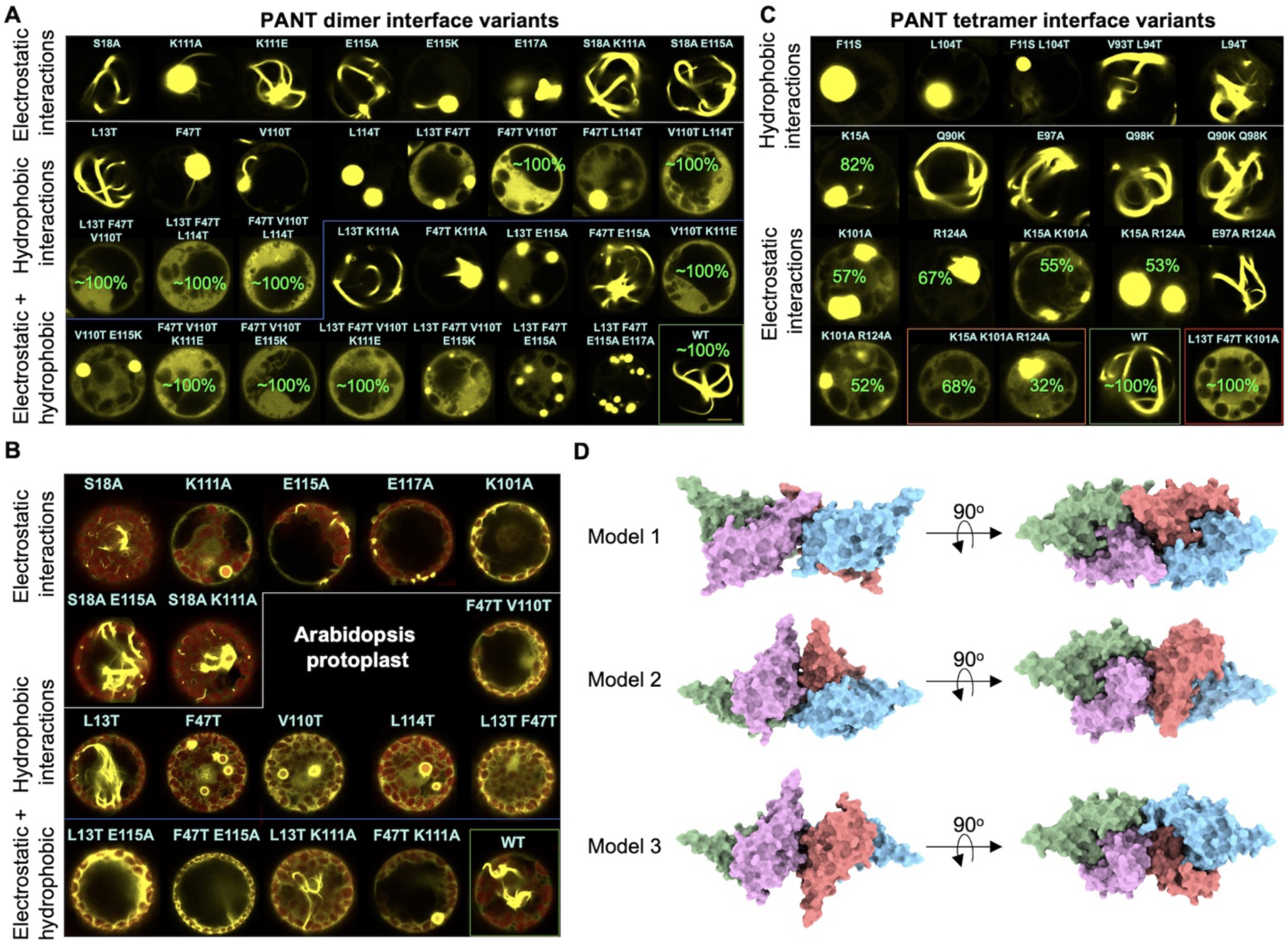
Mutational validation of AlphaFold2-predicted PANT dimer and tetramer interactions. (**A**) Subcellular localization of PANT dimer interface variants in rice protoplasts. Images were taken with the same confocal microscopy settings. (**B**) Subcellular localization of PANT dimer and tetramer interface variants in Arabidopsis protoplasts. Images were taken with the same confocal microscopy settings. (**C**) Subcellular localization of PANT tetramer interface variants in rice protoplasts. Images were taken with the same confocal microscopy settings. (**D**) PANT tetramers predicted with locally installed AlphaFold v2.2.4. Surface contacts of models 1-3 are shown.

**Fig. S8.**
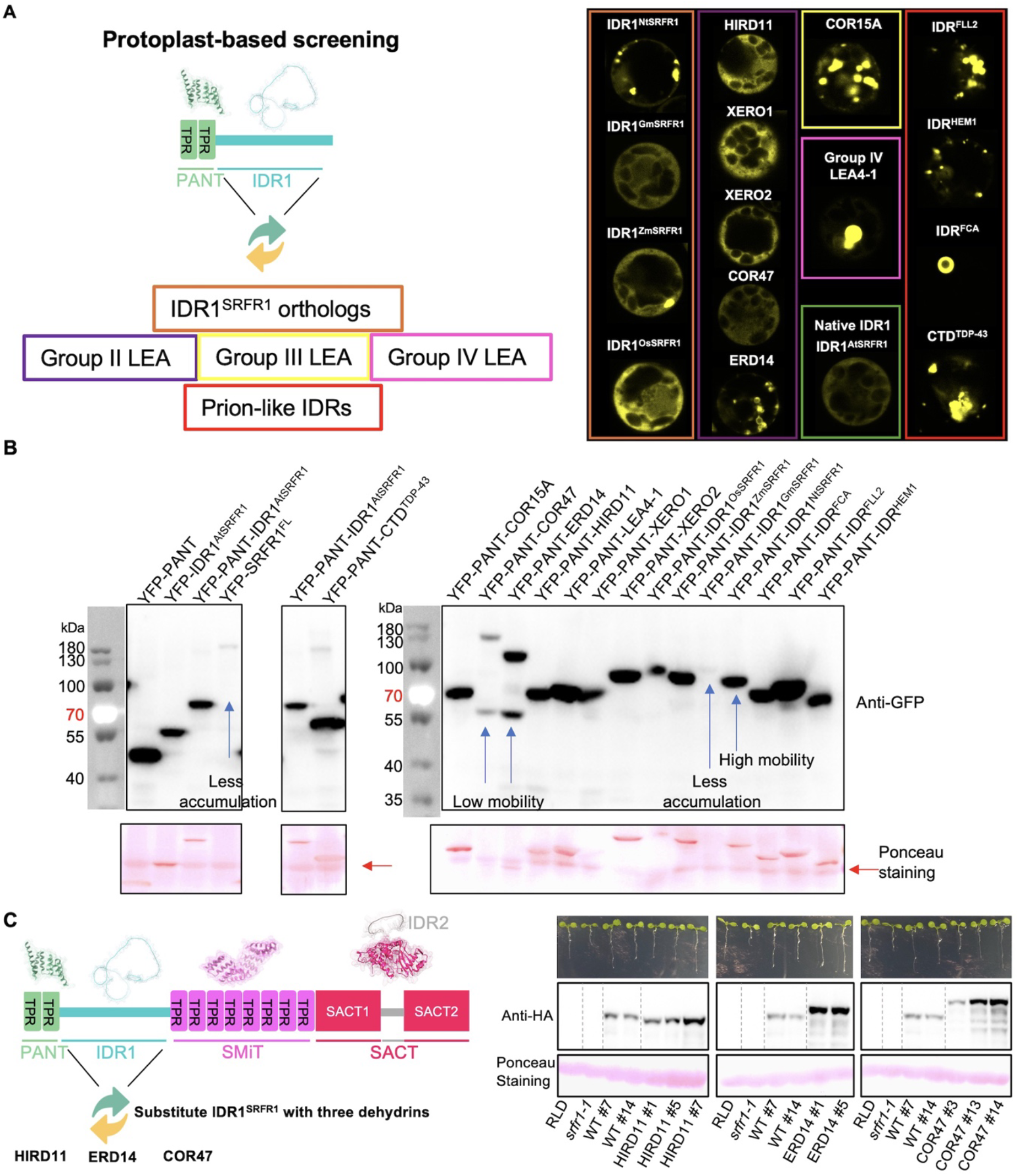
Functional substitution of IDR1^SRFR1^ with prion-like IDRs and LEA IDPs. (**A**) Schematic diagram of functional substitution strategy and subcellular localization of the PANT domain fused with IDR1 orthologs, group II, III, and IV LEA proteins, and prion-like IDRs from Arabidopsis and human proteins. Proteins are expressed in rice protoplasts. Images were taken with the same confocal microscopy settings. (**B**) Western-blot detection of YFP-tagged SRFR1 variants and PANT domain fused with IDR1 orthologs, plant LEA proteins and prion-like IDRs and expressed in rice protoplasts. Ponceau S staining of the Rubisco large subunit is shown as a protein loading control. (**C**) Primary root length and protein expression of YFP-HA-tagged SRFR1 with HIRD11, ERD14 or COR47 IRD1 substitutions. Representative images of 6-day-old seedlings are shown. Ponceau S staining of the Rubisco large subunit shows equal protein loading.

**Fig. S9.**
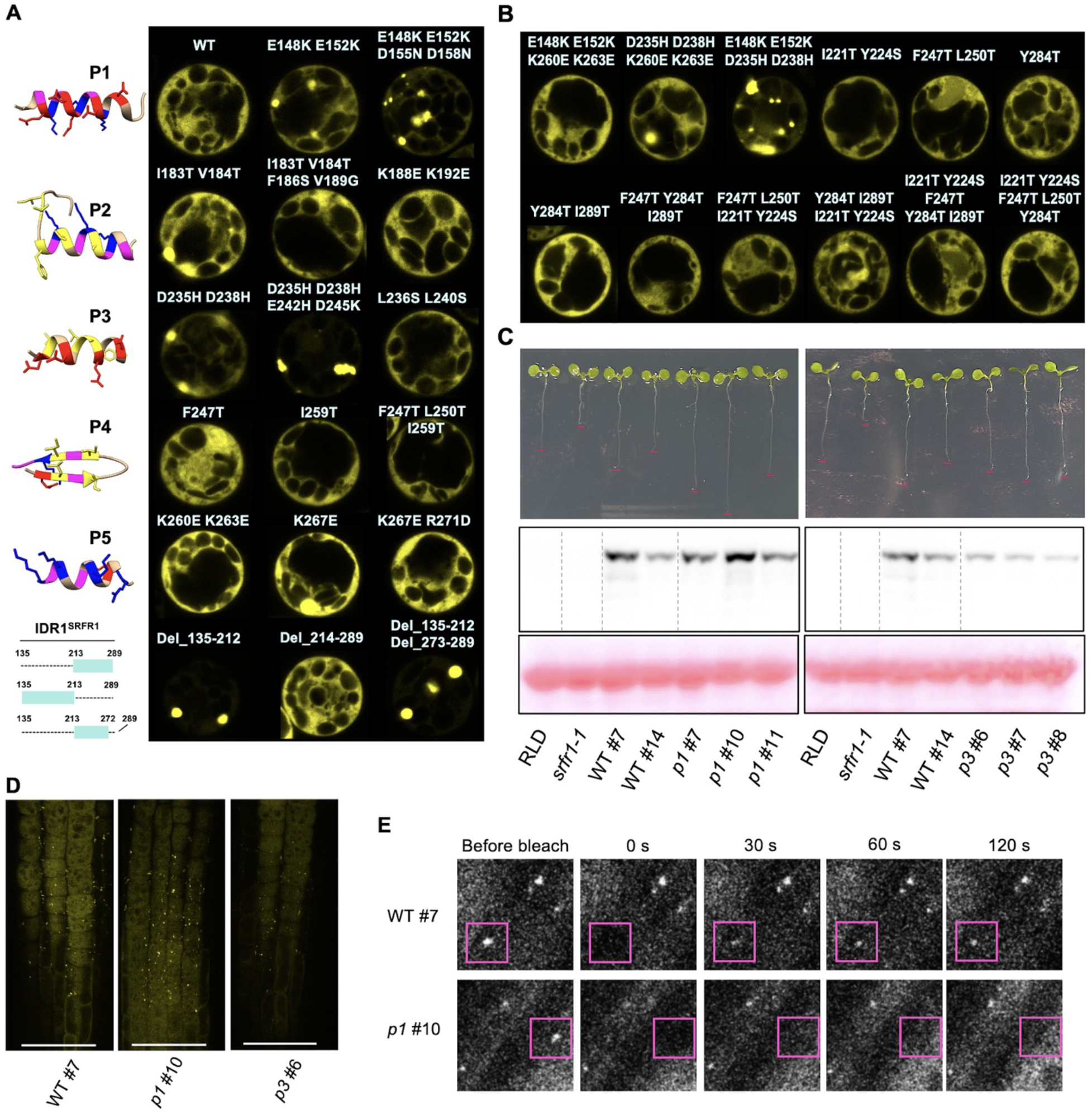
IDR1^SRFR1^ regulates SRFR1 biomolecular condensate formation and primary root growth. (**A** and **B**) Subcellular localization of PANT-IDR1^SRFR1^ variants in rice protoplasts. Potential functional segments are underlined and designated as P1 to P5. Structures of P1-P5 are predicted by PEP-FOLD3.5. Blue indicates positively charged amino acids; red indicates negatively charged amino acids; and yellow indicates hydrophobic amino acids. Images were taken with the same confocal microcopy settings. (**C**) Primary root length and protein expression of YFP-HA-tagged SRFR1 *p1* (E148K E152K D155N D158N) and *p3* (D235H D238H E242H D245K) variants. Ponceau S staining of the Rubisco large subunit shows equal protein loading. (**D**) Lateral root cap biomolecular condensate of YFP-HA-tagged SRFR1 *p1* (E148K E152K D155N D158N) and *p3* (D235H D238H E242H D245K) variants. Multiple single sections were taken with the same confocal microscopy settings; maximal projection images are shown, bar = 50 μm. (**E**) FRAP analysis of condensate fluidity was performed with 6-day-old seedlings; representative images are shown.

**Fig. S10.**
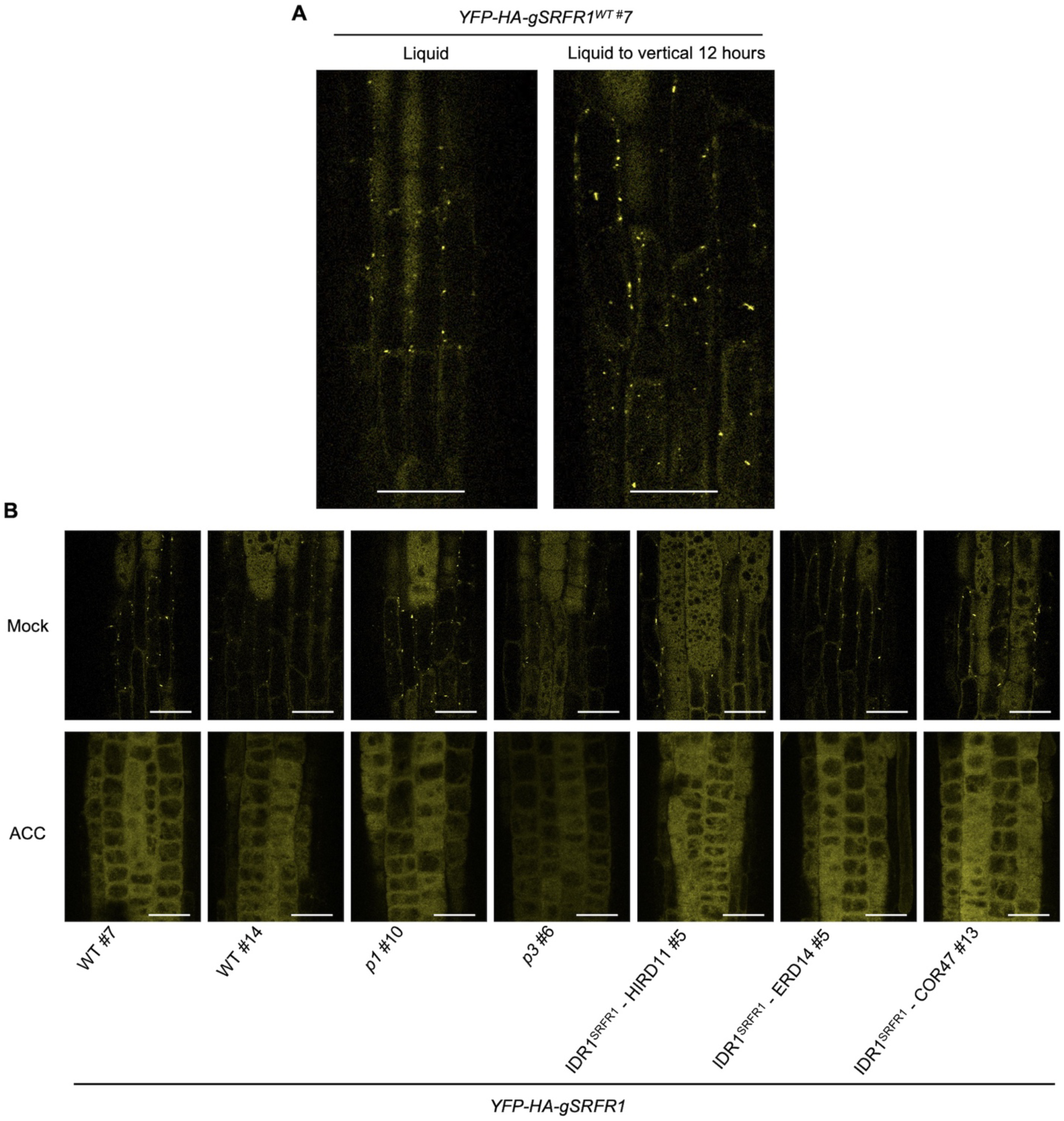
SRFR1 condensates are responsive to growth conditions and hormone treatment. (**A**) Lateral root cap SRFR1 condensates are induced after transferring from liquid medium to vertical growth. Images were taken with roots of 6-day-old seedlings 12 hours post transfer. Single surface section images were taken with the same confocal microscopy settings. (**B**) Lateral root cap biomolecular condensates formed by SRFR1 variants respond normally to ACC treatment. Images were taken with roots of 6-day-old seedlings 12 hours post ACC treatment. Top panel: single surface section images were taken with the same confocal microscopy settings. Bottom panel: multiple single sections were taken with the same confocal microscopy settings; maximal projection images showing the complete elimination of condensates; bar = 25 μm.

**Fig. S11.**
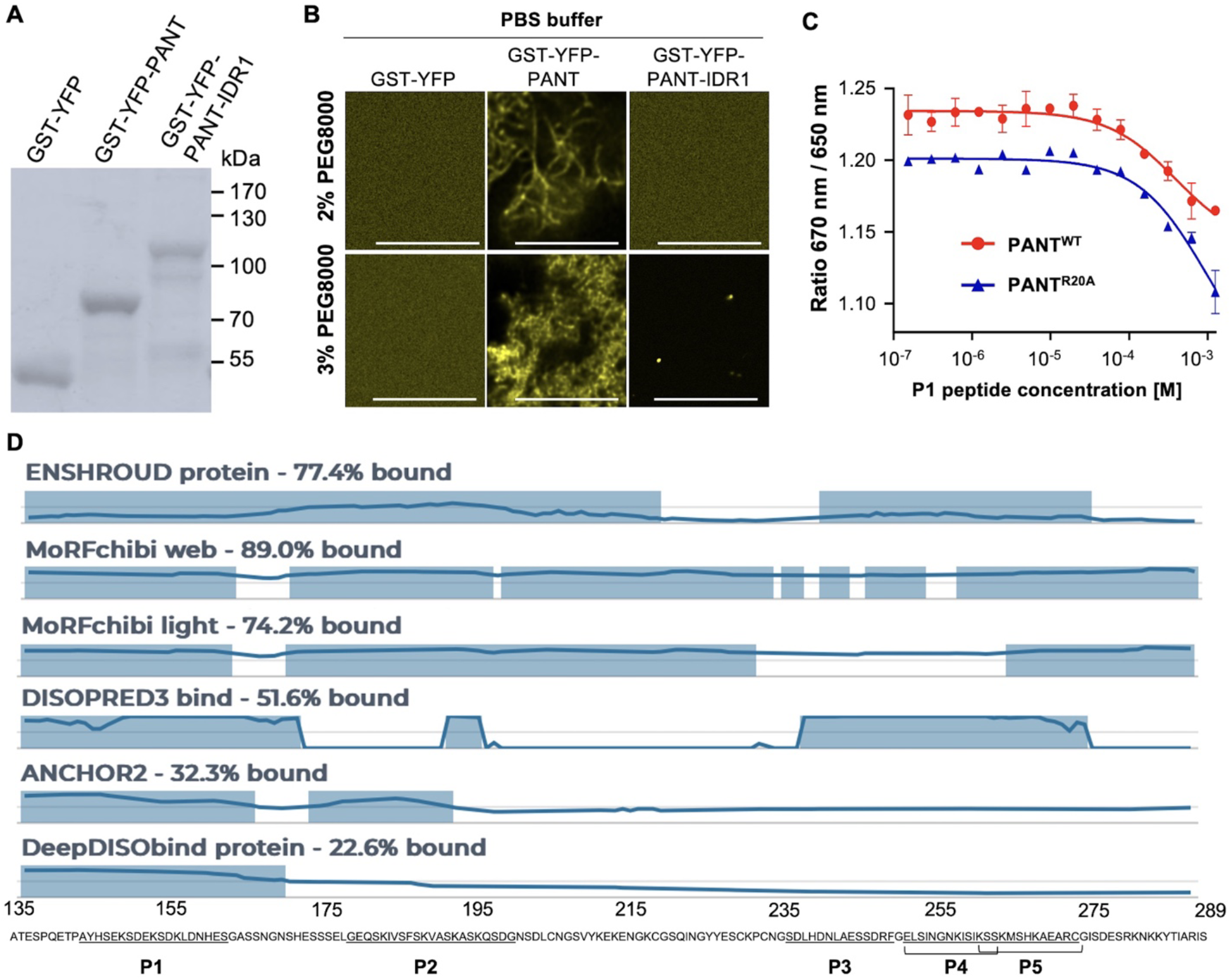
Interaction between PANT and IDR1^SRFR1^ regulate biomolecular condensate formation *in vitro*. (**A**) Coomassie blue staining of purified proteins. 2 mg protein was loaded for each sample. (**B**) Formation of PANT fibrils and PANT-IDR1^SRFR1^ condensate in PBS buffer induced by PEG8000. 30 μM protein was used for each. Images were taken 3 hours post PEG addition with the same confocal microscopy settings except for panels without fibril or condensate formation which were imaged with higher laser power and gain. Bar = 10 μm. (**C**) Quantification of PANT and P1^SRFR1^ binding affinity by microscale thermophoresis (MST). (**D**) Prediction of binding regions in IDR1^SRFR1^ by Critical Assessment of protein Intrinsic Disorder (CAID) prediction portal (*45, 48, 49, 71–74*).

**Fig. S12.**
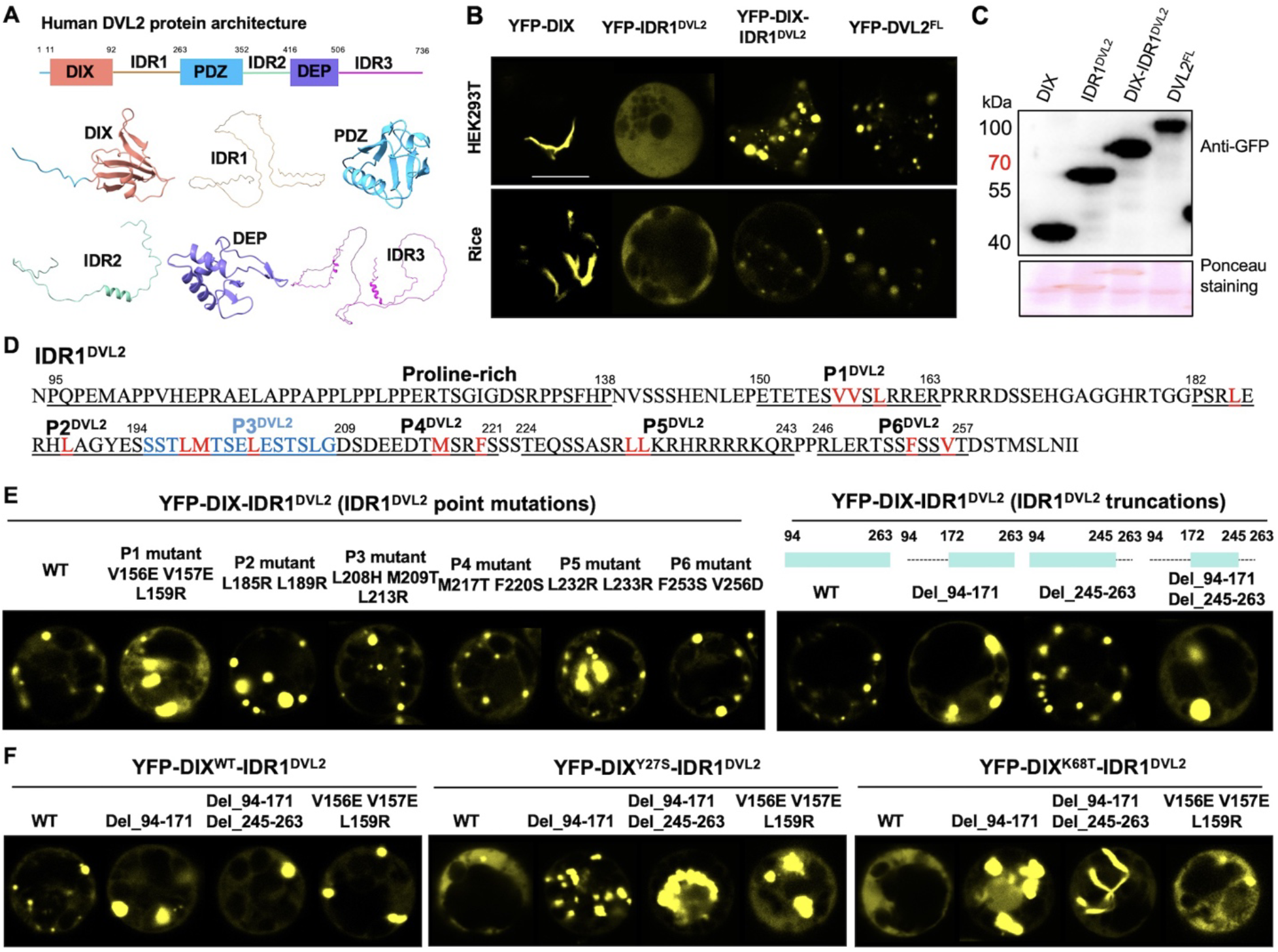
Interaction between DIX and IDR1^DVL2^ regulates DIX-IDR1^DVL2^ condensate formation. (**A**) Schematic diagram of DVL2 protein architecture. Structures of all domains are derived from AlphaFold2 prediction (AF-O14641-1). (**B**) Subcellular localization of DIX, IDR1^DVL2^, DIX-IDR1^DVL2^ and full-length DVL2 in human HEK293T cells and rice protoplasts. Images were taken with the same confocal microscopy settings. Bar = 10 μm. (**C**) Western-blot detection of DIX, IDR1^DVL2^, DIX-IDR1^DVL2^ and full-length DVL2 expression in rice protoplasts. Ponceau S staining of the Rubisco large subunit shows equal protein loading. (**D**) Potential functional segments in IDR1^DVL2^ are underlined and designated as P1^DVL2^ to P6^DVL2^. (**E** and **F**) Subcellular localization of DIX-IDR1^DVL2^ point mutation and truncation variants in rice protoplasts. Images were taken with the same confocal microscopy settings.

**Fig. S13.**
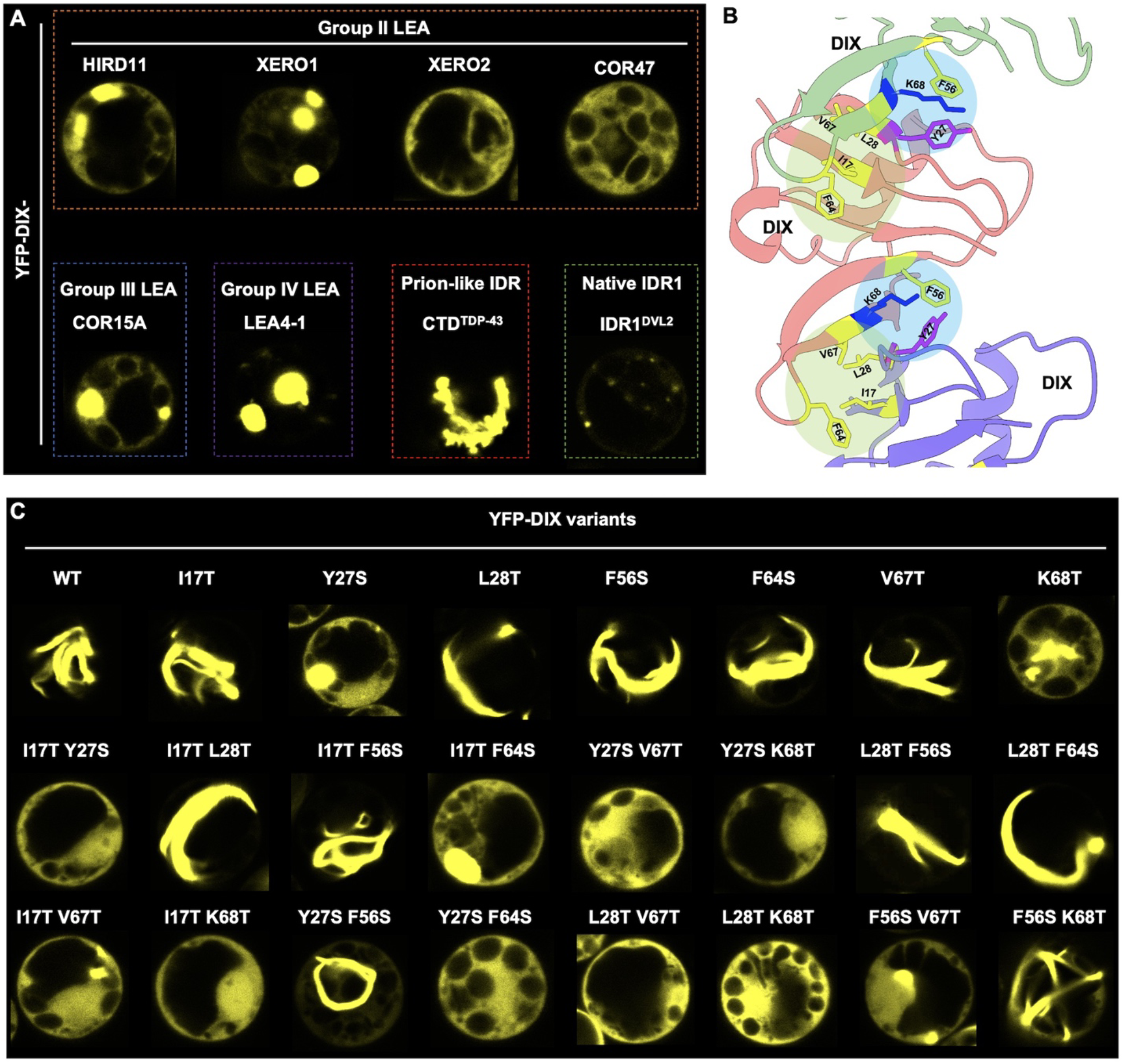
Regulation of DIX head-to-tail polymerization by IDR1^DVL2^. (**A**) Subcellular localization of the DIX domain fused with Arabidopsis group II, III and IV LEA proteins and IDR1 from human TDP-43 protein in rice protoplasts. Images were taken with same confocal microscopy settings. (**B**) Head-to-tail interaction interface of two DIX domain monomers. Structure was derived from PDB structure (6VCC) (*75*). (**C**) Subcellular localization of YFP-tagged DIX mutation variants in rice protoplasts. Images were taken with the same confocal microscopy settings.

